# *Ustilaginoidea virens* suppresses floral immunity through promoting GA biosynthesis by the effector SCRE9

**DOI:** 10.1101/2025.01.07.631775

**Authors:** Siwen Yu, Shaoqi Zhang, Xinhang Zheng, Xiaoai Li, Jiyang Wang, Guohua Duan, Shanshan Qiu, Dan Zhao, Nan Nan, Qianheng Yu, Chunquan Jiang, Zhao Peng, Dayong Li, Wenxian Sun

## Abstract

Rice false smut caused by *Ustilaginoidea virens* has become one of the most important rice diseases. *U. virens* specifically infects rice florets through stamen filaments. However, it is mysterious how *U. virens* inhibits floral immunity for successful colonization. Here, we show that a small cysteine-rich effector SCRE9 is a key virulence factor in *U. virens*. Expression of SCRE9 in rice not only suppresses immune responses and false smut resistance, but also significantly increases expression of expansin and gibberellin (GA) biosynthetic genes and GA content in young rice panicles. We further demonstrate that OsSIP1 (SCRE9-interacting protein 1) in rice is targeted by SCRE9 and interacts with the OsMADS63 and OsMADS68 transcription factors that transcriptionally regulate the GA synthesis gene *GA3ox1*. Interestingly, the chloroplast-localized OsSIP1 recruits OsMADS63/68 into the chloroplast. SCRE9 causes OsSIP1 to be translocated into the nucleus, thereby releasing OsMADS63/68 from the chloroplast into the nucleus to promote *GA3ox1* transcription. Therefore, SCRE9 promotes GA biosynthesis and expression of expansins in rice florets, thus loosening cell walls and overcoming the physical barrier during *U. virens* infection. The finding uncovers an unidentified floret infection strategy that offsets the deficiency in cell wall degrading ability in the unique floret colonizing fungus.

## Introduction

Plant pathogens secrete an array of effectors into the hosts to inhibit or interrupt defense signaling pathways for successful infection (Jones and Dangl 2006). Phytopathogenic fungi secrete apoplastic and intracellular effectors through conventional and unconventional ways (Tariqjaveed et al. 2021). Some apoplastic effectors, such as cell wall degrading enzymes, decompose plant cell walls and break the physical defensive barrier (Zhao et al. 2023; Sánchez-Torres et al. 2024). Intracellular effectors affect hormone signaling, alter protein stability, modulate gene transcription levels, mediate RNA degradation and alternative splicing to suppress plant immunity through targeting different immune components (Tariqjaveed et al. 2021). For instance, the *Puccinia striiformis* effector Pst18363 stabilizes the hydrolase TaNUDX23 and thereby inhibits reactive oxygen species (ROS) burst (Yang et al. 2019). The *Ustilago maydis* effector Cmu1 reduces the level of the salicylic acid (SA) precursor, thus inhibiting SA synthesis (Djamei et al. 2011). The *Verticillium dahliae* effector VdSCP41 inhibits SA signaling by interacting with transcription factors CBP60g and SARD1 to prevent the transcriptional activation of SA biosynthetic genes (Wang et al. 2011; Qin et al. 2018). The *P. striiformis* effector PNPi attenuates immune responses by inhibiting the interaction of the TGA-bZIP transcription factor TGA2.2 with NPR1, a key regulatory protein in SA signaling (Wang et al. 2016). Besides, the *Xanthomonas* effector kinase XopC2 promotes JAZ protein degradation and enhances JA signaling, thereby facilitating stomatal opening and bacterial infection (Wang et al. 2021). In contrast to many effectors that target SA and JA signaling pathways, no fungal effector has been identified to regulate gibberellin (GA) signaling pathway to inhibit plant immunity.

GA and its biosynthetic genes regulate plant growth and immunity. GA often interferes with SA- and JA-mediated defense pathways by affecting DELLA stability (De Vleesschauwer et al. 2013; De Vleesschauwer et al. 2016; Samanta and Roychoudhury 2024; Wan et al. 2024). Exogenous GA reduces rice resistance to the (semi-) biotrophic pathogens *M. oryzae* and *X. oryzae* pv. *oryzae* (De Vleesschauwer et al. 2013). GA20 oxidases (GA20ox) catalyze successive oxidation steps and promote GA biosynthesis (Hedden and Thomas 2012). *OsGA20ox3*-silencing rice plants exhibit a semi-dwarf phenotype and increased resistance to *M. oryzae* and *X. oryzae* pv. *oryzae*, whereas the *OsGA20ox3-*overexpressing transgenic plants have an elevated content of active gibberellins and show a slender phenotype and increased susceptibility to rice pathogens (Qin et al. 2013).

Through promoting cell proliferation and expansion, GA regulates plant growth and development, such as stem elongation, leaf expansion, especially flower organ development (Gao et al. 2017). GA can induce expansin expression (Lee and Kende 2001; Azeez et al. 2010; Sun et al. 2011; Li et al. 2019). Expansins unlock the network of wall polysaccharides, permitting cell wall loosening (Cosgrove 2000; Cosgrove 2024). In gladiolus flowers, the expansin GgEXPA1 is involved in cell elongation in stamen filament and pistil style (Azeez et al. 2010). Cell wall loosening caused by expansins is conducive to pathogen infection. For instance, *X. oryzae* pv. *oryzae* infection induces expression of expansin genes and causes cell wall loosening, thereby rendering host rice vulnerable to the pathogen (Ding et al. 2008).

*Ustilaginoidea virens* causes one of the most devastating diseases in rice and poses a severe threat to rice commercial production worldwide (Sun et al. 2020). *U. virens* produces a great amount of mycotoxins, such as ustiloxins, ustilaginoidins, and sorbicillinoids, which threaten human and animal health (Yu et al. 2023). Interestingly, *U. virens* has its unique infection mode, exclusively infecting stamen filaments (Hu et al. 2013). In addition, no evident specialized feeding structure (e.g. haustoria) or appressorium has been observed during *U. virens* infection. According to microscopic observation, the arrangement of microfilaments in cell wall of stamen filaments is extremely loose. Because *U. virens* encodes many fewer cell wall degradation enzymes than other phytopathogenic fungi, loose cell wall structure in stamen filament might be Achilles’ Heel in rice for *U. virens* infection (Hu et al. 2013; Zhang et al. 2014). However, it is still mysterious how *U. virens* breaks physical barrier of floral cell walls and whether GA-promoted cell wall loosening plays an important role in *U. virens* infection.

*U. virens* effectors have been identified to inhibit rice immunity through various strategies. The effector SCRE1 outcompetes SnRK1a and ATP for XB24 binding, thereby inhibiting SnRK1a-mediated XB24 phosphorylation and ATP hydrolysis (Yang et al. 2022). The nuclear effector SCRE4 inhibits the expression of *OsARF17* encoding a positive immune regulator (Qiu et al. 2022). Notably, SCRE6 is the first identified effector phosphatase in fungi. The effector specifically dephosphorylates and stabilizes the negative immune regulator OsMPK6, thus promoting *U. virens* infection (Zheng et al. 2022). The UvPr1a effector directly degrades OsSGT1, a positive regulator of immunity against rice pathogens (Chen et al. 2022b). A couple of *U. virens* effectors inhibit rice immunity through regulating epigenetics. UvSec117 interacts with and recruits the histone deacetylase OsHDA701 into host nuclei, which mediates H3K9 deacetylation and inhibits expression of immune-related genes (Chen et al. 2022a). Besides, Uv1809 targets the histone deacetylase OsSRT2 and enhances histone deacetylation, thereby reducing H4K5ac and H4K8ac levels and suppressing defense gene expression (Chen et al. 2023). A multifunctional effector UvCBP1 can block chitin-induced immunity (Li et al. 2021). Besides, UvCBP1 interacts with rice scaffold protein OsRACK1A and outcompetes its interaction with NADPH oxidase OsRBOHB, thus inhibiting ROS generation (Li et al. 2022). Interestingly, several effectors, such as SGP1 and UvGHF1, not only function as essential virulence factors during *U. virens* infection, but also are recognized as microbe-associated molecular patterns (Song et al. 2021; Zou et al. 2023). Although the increasing number of *U. virens* effectors have been functionally identified, the mechanism how *U. virens* effectors specifically target rice floral immunity remains unclear.

In this study, we show that a small cysteine-rich effector 9 (SCRE9) is required for *U. virens* virulence. Furthermore, we find that SCRE9 targets OsSIP1, which inhibits GA biosynthesis through confining the transcription factors OsMADS63/68 to the chloroplast. However, SCRE9 induces the biosynthesis of endogenous gibberellins in rice panicles through releasing the chloroplast-confined MADS transcription factors. Consistently, SCRE9 promotes the expression of expansins in rice young panicles to inhibit floral defenses and therefore facilitates *U. virens* infection. The findings reveal an unidentified floret infection mechanism that facilitates *U. virens* floral colonization.

## Results

### SCRE9 is a virulence effector in *U. virens*

SCRE9 (UV8b_02533) is predicted to be a secreted cysteine-rich effector with 142 amino-acid residues in *U. virens*. SCRE9 contains a putative signal peptide (SP) at its N-terminus (Supplementary Fig. S1, A and B). We performed a yeast secretion assay to validate the functionality of the putative SP of SCRE9 (Fang et al., 2019). The SP-encoding sequences of *SCRE9* and *Avr1b* were fused in frame with a *SUC2* gene variant that encodes truncated invertase lacking its own signal peptide. These constructs were then transformed into the invertase deficient YTK12 strain. The culture medium of the yeast strains transformed with *Avr1b^SP^-SUC2* and *SCRE9^SP^-SUC2* turned red after the addition of 2,3,5-triphenyltetrazolium chloride (TTC) (Fig. 1A). However, the culture medium of non-transformed YTK12 and *Mg87^N25^-SUC2* transformed strains, as negative controls, was colorless. These data indicate that the putative signal peptide of SCRE9 is able to guide protein secretion. Next, to determine whether SCRE9 is translocated into plant cells during infection, the *M. oryzae* transformants ectopically expressing SCRE9-GFP were inoculated into rice sheaths (Qiu et al. 2022; Zheng et al. 2022). Green fluorescence was observed in the biotrophic interfacial complex (BIC, plant-derived membrane-rich structure) at 30- to 36-h after inoculation. The results indicate that SCRE9-GFP expressed in *M. oryzae* is secreted into rice cells via BICs during infection. *M. oryzae* strains expressing Avr-Pia-GFP and GFP were inoculated into rice sheaths as positive and negative controls, respectively. Green fluorescence from Avr-Pia-GFP was observed in BICs, whereas GFP fluorescence was only visible in invasive hyphae of the GFP-expressing strain (Supplementary Fig. S1C). These results suggest that SCRE9 is translocated into plant cells during fungal infection. Furthermore, the expression pattern of *SCRE9* was examined during *U. virens* infection.Reverse transcription quantitative PCR (RT-qPCR) assays showed that *SCRE9* expression was dramatically induced during *U. virens* infection with two peaks at 4 d and 8 d after inoculation (Supplementary Fig. S1D), suggesting that SCRE9 might play an important role in *U. virens* infection.

**Figure 1.**
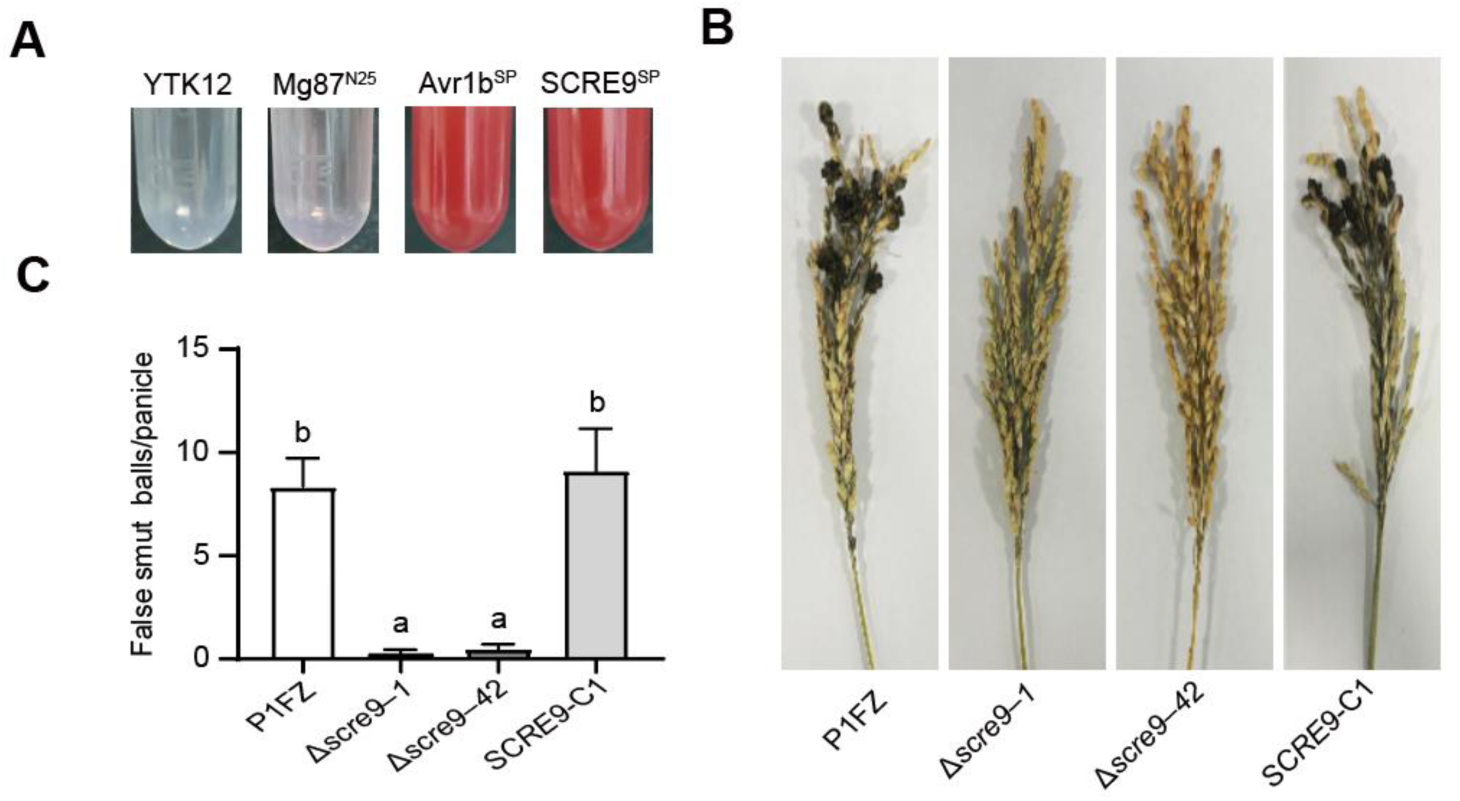
SCRE9 is a virulence effector in *Ustilaginoidea virens*. A, The putative signal peptide of SCRE9 guided the secretion of the invertase SUC2. The culture medium of the SCRE9^SP^-SUC2 expressing yeast strain turned red after the addition of TTC. The YTK12 and Mg87^N25^-SUC2 expressing strains were used as negative controls, while the Avr1b^SP^-SUC2 expressing strain was used as a positive control. B-C, Disease symptoms and the average number of diseased grains per inoculated panicle after inoculation with different *U. virens* strains. The wild-type (P1FZ), Δ*scre9* knockout mutant (Δ*scre9*-1 and Δ*scre9*-42) and complemented (*SCRE9−C1*) strains were injection-inoculated into young panicles of the susceptible rice cultivar JN853. The images were captured and diseased grains were counted at one month after inoculation. Data are presented as mean ± standard error (SE) (*n* = 30). Different letters (a vs. b) indicate significant difference in the number of diseased grains (One way ANOVA followed by Duncan’s multiple range test).

To confirm the virulence function of SCRE9, the *SCRE9* gene replacement mutants were generated through CRISPR/Cas9-induced homologous recombination and were then confirmed via Southern blot analyses (Supplementary Fig. S2A). The complemented strains were constructed by introducing the *SCRE9* gene with the strong promoter of *UVR_08266* into the mutant strain. Expression of SCRE9-FLAG in complemented strains was detected via immunoblotting (Supplementary Fig. S2B). Subsequently, the wild-type, Δ*scre9* and Δ*scre9-C* complemented strains were injection-inoculated into rice panicles of the susceptible cultivar JN853. Diseased grains on rice panicles inoculated with the wild-type and complemented strains were significantly more than those on the panicles inoculated with the Δ*scre9* strains (Fig. 1, B and C). These results indicate that SCRE9 plays an essential role in *U. virens* virulence to rice.

### SCRE9 suppresses pattern-triggered immunity and promotes rice growth

To investigate whether SCRE9 suppresses rice immunity, the IE383, IE409 and IE421 transgenic lines with dexamethasone (DEX)-induced expression of SCRE9 were generated and were then confirmed through immunoblotting for subsequent studies (Supplementary Fig. S3A). Both chitin and flg22 triggered a rapid and robust generation of oxidative burst in the wild-type and IE383 transgenic rice plants without DEX induction. By contrast, the induced oxidative burst was significantly suppressed in the IE383 transgenic line after DEX treatment (Fig. 2, A and B; Supplementary Fig. S3, B and C). Furthermore, RT-qPCR assays showed that expression of *OsPR10* and *OsPR1b* was strongly up-regulated by flg22 and chitin in the wild-type and IE383 transgenic seedlings without DEX treatment. By contrast, flg22-and chitin-induced expression of these defense-related genes was significantly suppressed in the DEX-treated IE383 transgenic line (Fig. 2, C to E). These results indicate that SCRE9 inhibits pattern-triggered immunity in rice.

**Figure 2.**
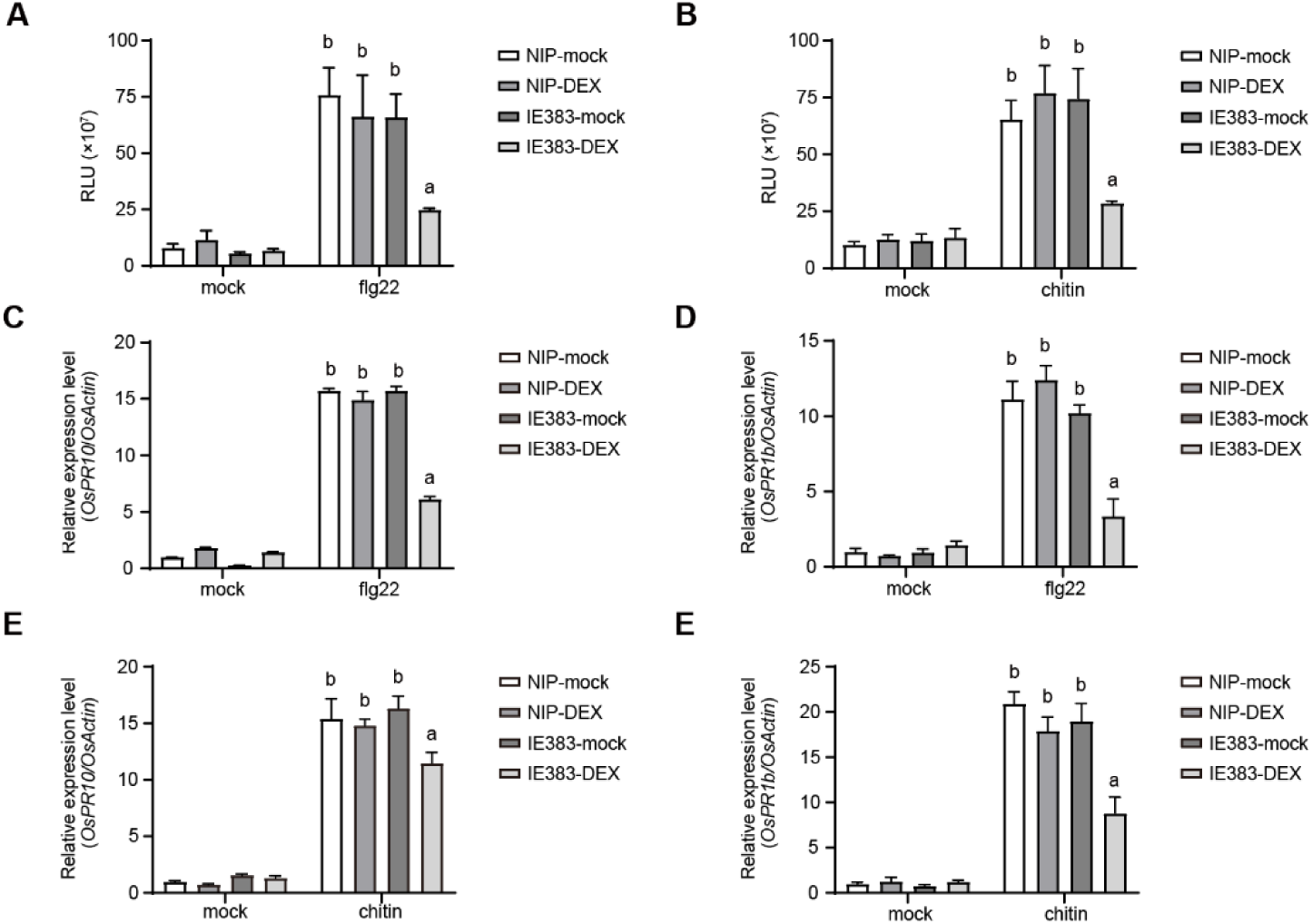
DEX-induced expression of SCRE9 inhibits immune responses in rice. A-B, flg22- and chitin-triggered ROS burst in the wild-type and IE383 transgenic lines with or without DEX treatment. The wild-type and transgenic seedlings were treated with DEX (10 µM) or Mock solution for 24 h followed by flg22 (1 µM), chitin (10 µg/mL) or mock treatment. ROS burst was detected immediately after PAMP treatments for 24 min. The peak areas in Figure S3B-C were calculated using ImageJ and are shown as mean ± SE (*n* = 3). RLU, relative luminescence unit. C-F, The flg22- (C, D) and chitin-(E, F) induced expression of *OsPR10* (C, E) and *OsPR1b* (D, F) in the wild-type and transgenic lines with DEX or mock treatment. The wild-type and IE383 transgenic seedlings were treated with DEX (10 µM) or mock solution for 24 h followed by flg22 (1 µM) or chitin (10 µg/mL) treatment. Expression of *OsPR10* and *OsPR1b* was detected at 6 h after flg22 and chitin treatments by RT-qPCR and was normalized to the reference gene *OsActin*. Different letters above error bars indicate significant differences in the *OsPR10* and *OsPR1b* transcript levels in the wild-type and transgenic lines under different treatments (One way ANOVA followed by Duncan’s multiple range test). Representative data from three independent experiments are presented as mean ± SE (*n* = 3).

Meanwhile, we generated and confirmed the CE6 and CE13 transgenic lines with constitutive expression of SCRE9 through immunoblotting (Supplementary Fig. S3A). The effects of SCRE9 expression on rice growth and development were then examined. The CE6 and CE13 transgenic lines showed more robust growth but exhibited no significant difference in tillering, grains per panicle, seed setting rate and 1000-grain weight (Supplementary Fig. S4). Taken together, these results suggest that SCRE9 promotes plant growth but suppresses immune responses in rice.

### SCRE9 induces expression of the genes encoding expansins, pollen allergens and β-glucosidases

To illustrate molecular mechanisms of SCRE9 inhibiting rice immunity, RNA-seq analyses were conducted to compare gene expression profilings in the wild-type and IE383 transgenic rice panicles after DEX treatment. Principal component analysis (PCA) showed that RNA-seq data from the wild-type and transgenic lines were clearly separated in two distinct groups, indicating that the wild-type and transgenic lines exhibit significant difference in genome-wide gene expression (Fig. 3A). Transcriptome analyses revealed 761 differentially expressed genes (DEGs) with∣ Log2 (fold change)∣ ≥ 1 and *P* < 0.05, among which 583 genes were up-regulated and 178 genes were down-regulated (Fig. 3, Band C). GO enrichment analysis found that DEGs were enriched in such terms as plant-type cell wall organization (GO:0009664), hydrolase activity, acting on glycosyl bonds (GO:0016798), plant-type cell wall organization or biogenesis (GO:0071669), external encapsulating structure organization (GO:0045229), intramolecular lyase activity (GO:0016872), hydrolase activity, hydrolyzing O-glycosyl compounds (GO:0016798), and cell wall organization (GO:0071555) (Fig. 3D).

**Figure 3.**
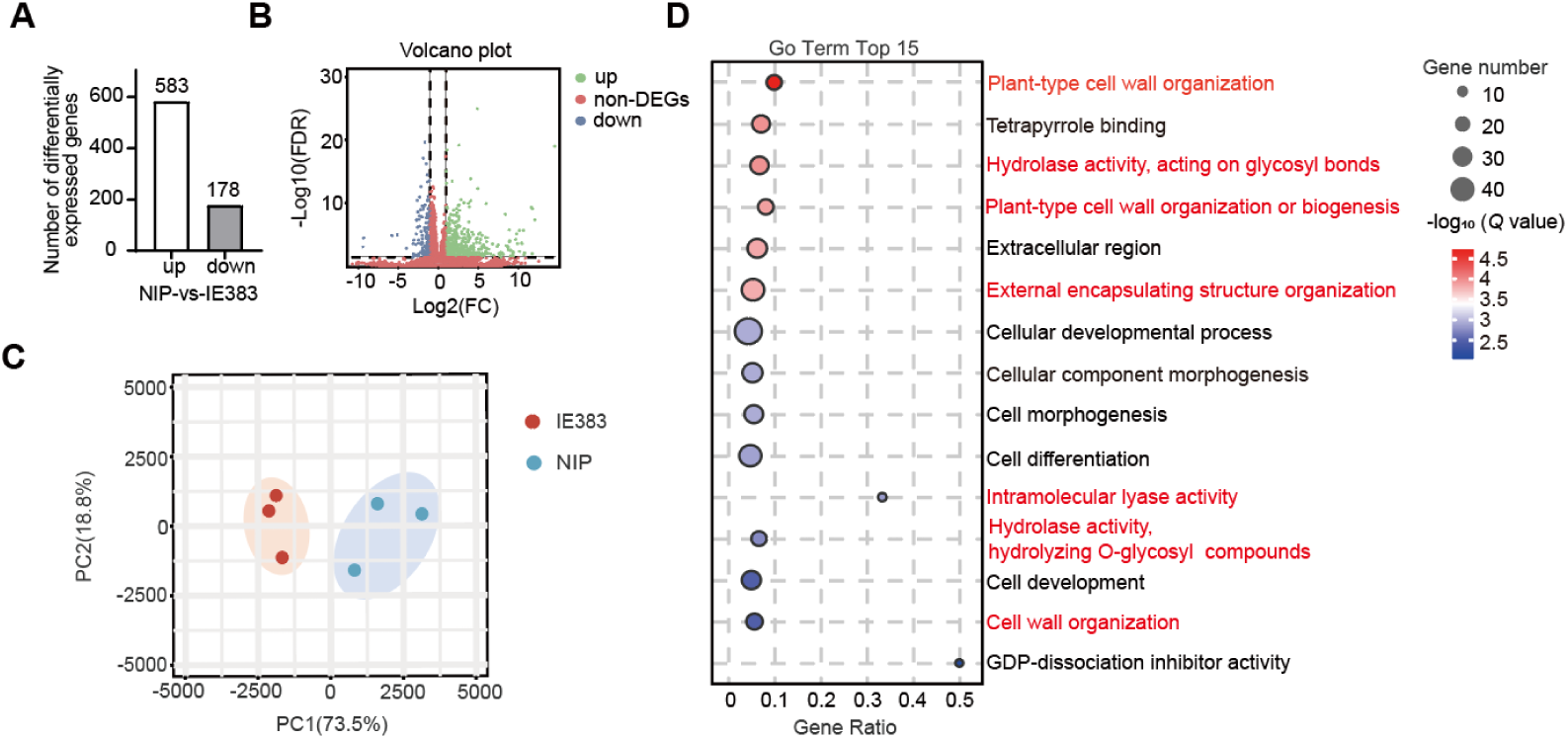
Differentially expressed genes are enriched in cell wall organization-associated terms between the wild-type and SCRE9-expressing transgenic rice lines. A, The number of differentially expressed genes (DEGs) between the wild-type and SCRE9-expressing IE383 transgenic rice lines. B, Volcano plot to show DEGs between the wild-type and IE383 transgenic lines. Green and blue colors represent up-regulated and down-regulated genes, respectively, and red represents non-DEGs. The genes with ∣ Log2(fold change) ∣ ≥ 1 and *P* < 0.05 were identified as DEGs. FDR, false discovery rate; FC, fold change. C, Principal component analysis (PCA) to show the reproducibility and reliability of transcriptome data. Gene expression profiles from three samples of the IE383 transgenic line were grouped as a clade, but were separated from the wild-type (NIP) samples. D, GO enrichment analysis revealed the top 15 GO terms enriched in DEGs between the wild-type and transgenic rice lines. The size of circles represents the number of DEGs mapped to specific GO terms. The color bar indicates *Q* values. Red color represents the GO terms associated with cell wall organization and degradation.

Plant cell wall is the major physical barrier for fungal infection. Compared with many other plant pathogenic fungi, *U. virens* has a reduced ability to break plant cell walls (Zhang et al., 2014). Notably, many genes encoding cell wall loosening-associated proteins, such as expansins, pollen allergens and β-glucosidases, were dramatically up-regulated in SCRE9-expressing transgenic panicles (Supplementary Table S1). Previous studies have shown that stamen filaments are the initial infection sites of *U. virens* (Tang et al. 2013). Expansins and pollen allergens are involved in loosening cell wall of stamen filaments and stigma (Cosgrove 2000; Azeez et al. 2010; Cosgrove 2024). Meanwhile, β-glucosidases are a class of glycoside hydrolases that degrade cellulose in cell walls. Therefore, expansins, pollen allergens, and β-glucosidases might contribute to cell wall rearrangement in rice floral organs, which is conducive to the invasion of *U. virens*. Taken together, induced expression of these genes suggests that *U. virens* might promote cell wall degradation and expansion for successful infection through the action of SCRE9.

### SCRE9 up-regulates GA biosynthesis related genes and GA accumulation

KEGG pathway enrichment analysis was further performed to understand how SCRE9 regulates rice defense and growth. The enriched KEGG pathways for DEGs included phenylpropanoid biosynthesis (ko00940), biosynthesis of secondary metabolites (ko01110), metabolic pathways (ko01100), diterpenoid biosynthesis (ko00904), starch and sucrose metabolism (ko00500), pentose and glucuronate interconversions (ko00040), endocytosis (ko04144), and so on (Fig. 4A). Notably, diterpenoid is a precursor of gibberellin. In the diterpenoid biosynthesis pathway (ko00904), multiple GA biosynthetic genes were up-regulated in SCRE9-expressing transgenic plants as revealed by transcriptome data (Fig. 4B). Subsequently, differential expression of these GA biosynthetic genes was confirmed by RT-qPCR (Fig. 4C).

**Figure 4.**
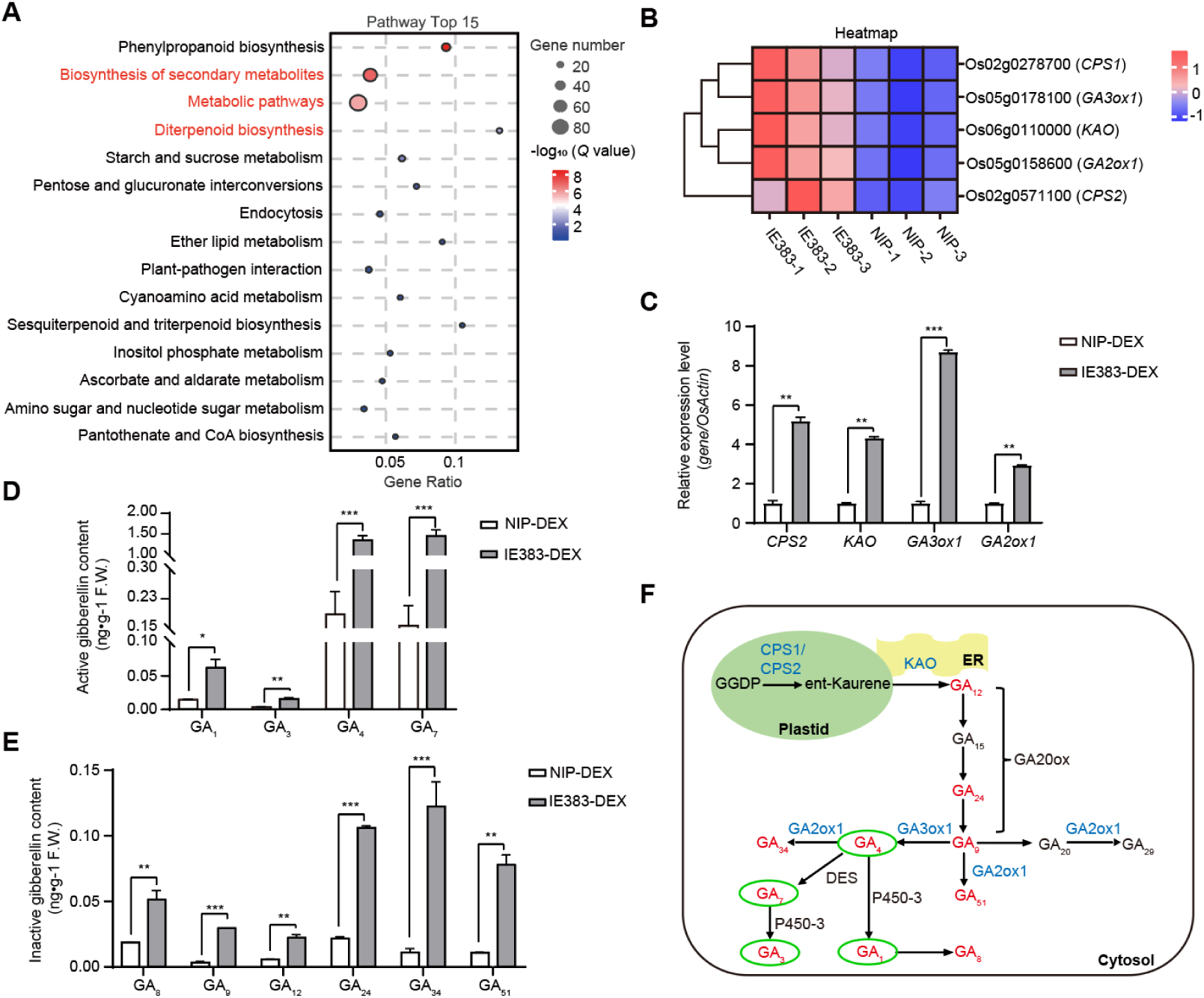
Ectopic expression of SCRE9 transcriptionally up-regulates GA biosynthesis-related genes and elevates GA accumulation in rice panicles. A, The top 15 enriched KEGG pathways for DEGs as revealed by KEGG pathway enrichment analysis. B, A heatmap to show the expression profiles of gibberellin biosynthesis-related genes in the wild-type and SCRE9-expressing transgenic rice panicles. C, The expression levels of gibberellin biosynthesis-related genes in the wild-type and transgenic rice panicles as detected by qRT-PCR. Representative data from three independent experiments are presented as mean ± SE (*n* = 3). D-E, The content of gibberellins in the wild-type and SCRE9-expressing transgenic rice panicles as determined by HPLC tandem mass spectrometry. GA_1_, GA_3_, GA_4_, and GA_7_ are active gibberellins (D). GA_8_, GA_9_, GA_12_, GA_24_, GA_34_, and GA_51_ are inactive gibberellins or intermediates (E). Representative data from three independent experiments are shown as mean ± SE (*n* = 3). In C-E, the wild-type and transgenic lines were treated with DEX (10 μM) for 24 h. *, *P* < 0.05; **, *P* < 0.01; ***, *P* < 0.001. F, A schematic diagram of gene expression profiles in the gibberellin biosynthesis pathway. GGDP, geranyl geranyl diphosphate; ER, endoplasmic reticulum. Red color indicates the type of gibberellins with increased accumulation, while blue represents up-regulated genes in the SCRE9-expressing transgenic line. Black color indicates the type of gibberellins with no change and non-DEGs; green circles indicate active gibberellin.

Next, the content of gibberellins was quantified in rice panicles of the wild-type and IE383 transgenic plants after DEX treatment. The results showed that the transgenic rice panicles produced significantly more active gibberellins including GA1, GA3, GA4 and GA7 than did the wild-type rice panicles. Particularly, the abundance of the major types of active GAs, GA4 and GA7, in rice panicles was much higher in the transgenic line compared with the wild type (Fig. 4D). Meanwhile, the content of inactive gibberellins and intermediates including GA8, GA9, GA12, GA24, GA34, and GA51 was also significantly higher in the transgenic lines compared with the wild type (Fig. 4E). The data indicate that expression of SCRE9 in transgenic rice line transcriptionally upregulates GA biosynthesis-related genes and thereby significantly increases the content of gibberellins in rice panicles (Fig. 4F).

### SCRE9 inhibits rice immunity through regulating GA signaling

To investigate the effect of GAs on regulating disease susceptibility, rice plants were treated with GA4+7, a GA biosynthesis inhibitor paclobutrazol (PAC), and mock control followed by pathogen inoculation. *U. virens* inoculation assays showed that the wild-type plants generated significantly more diseased grains on the inoculated panicles after GA4+7 treatment compared with mock control, whereas the PAC-treated panicles produced fewer diseased grains after inoculation (Fig. 5, A and B). In addition, DEX-induced expression of SCRE9 caused the transgenic lines to generate more diseased grains. The wild-type plants showed no difference in false smut resistance regardless of DEX treatment. After simultaneous treatment with GA and DEX, susceptibility to false smut disease in the *SCRE9* transgenic rice plants was further increased (Fig. 5 A and B). However, the DEX-treated IE383 transgenic line exhibited an elevated resistance to *U. virens* infection after PAC treatment, which was as strong as that in the PAC-treated wild-type plants. These data indicate that ectopic expression of SCRE9 and exogenous GAs both cause rice plants more susceptible to rice false smut.

**Figure 5.**
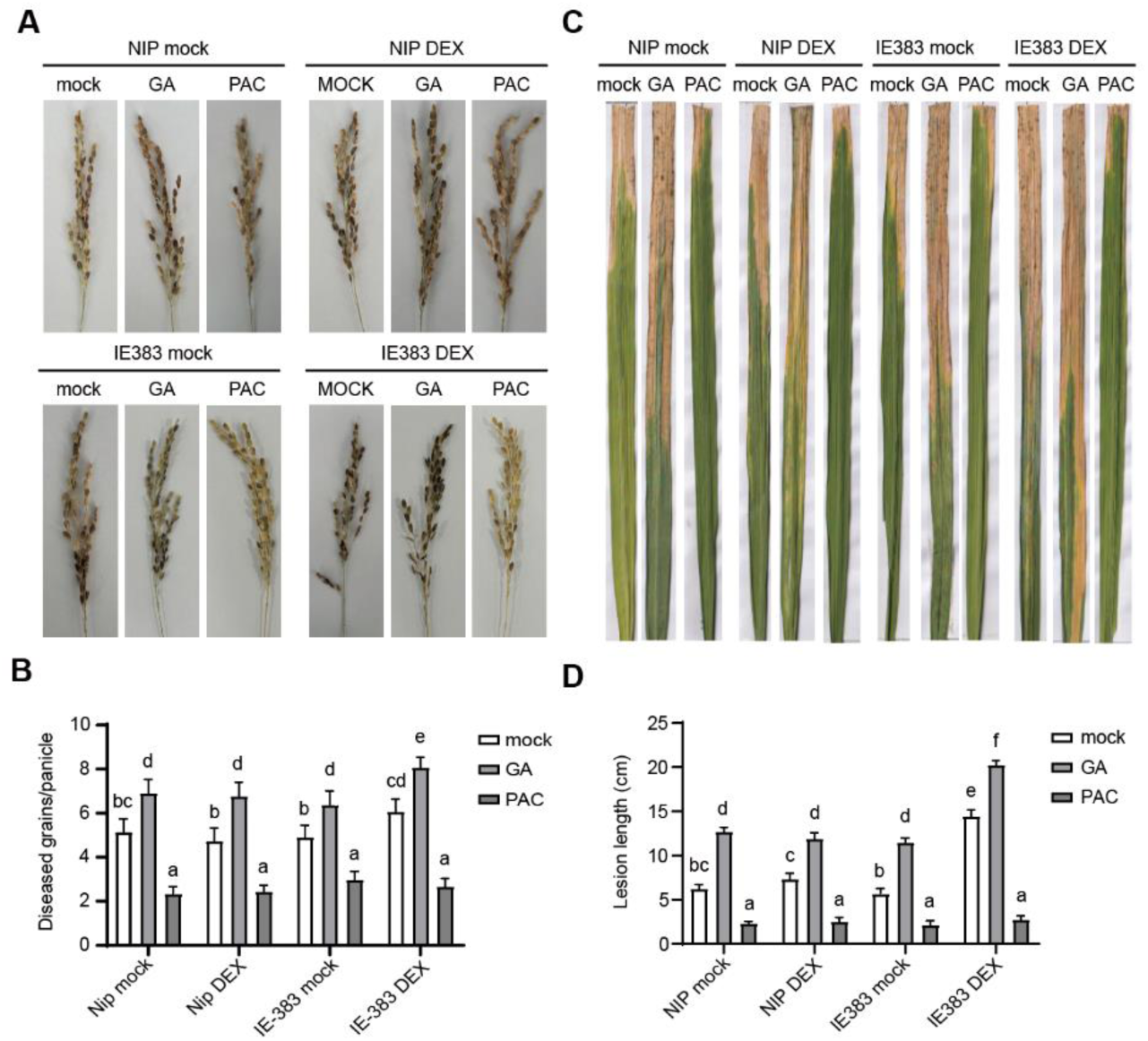
SCRE9 and GA attenuate disease resistance in rice. A-B, Disease symptoms and diseased grains on inoculated panicles of the wild-type and SCRE9-expressing transgenic rice lines after *U. virens* inoculation. Rice plants were sprayed with GA_4+7_ (60 μM) and PAC (20 μM) once a week for about one month before inoculation. Rice panicles were then treated with DEX (10 μM) or mock control for 24 h followed by injection inoculation with *U. virens*. The diseased grains were counted at about one month after inoculation. Representative data from three independent assays are presented as mean ± SE (*n* = 30). C-D, Disease symptoms (C) and disease lesion lengths (D) on inoculated leaves of the wild-type and SCRE9-expressing transgenic plants after *X. oryzae* pv. *oryzae* infection. Rice plants were pretreated with GA_4+7_ and PAC before bacterial inoculation as described above. Rice leaves were treated with DEX (10 μM) or Mock control for 24 h followed by clipping inoculation with *X. oryzae* pv. *oryzae*. The images were captured and disease lesion lengths were measured at 15 days post-inoculation (dpi). Representative data from three independent assays are presented as mean ± SE (*n* = 10). Different letters indicate significant difference in the number of disease grains (in B) and disease lesion length (in D) among different treatments (One way ANOVA followed by Duncan’s multiple range test).

Furthermore, we showed that *X. oryzae* pv. *oryzae* caused longer disease lesions on GA-treated rice leaves, but produced shorter disease lesions on PAC-treated leaves compared with mock control (Fig. 5, C and D). Likewise, the IE383 transgenic rice plants exhibited longer disease lesions on the *Xoo*-inoculated leaves than did the wild-type plants after DEX treatment. The wild-type plants showed similar length of disease lesions on the *Xoo*-inoculated leaves regardless of DEX treatment. The data indicate that SCRE9 enhances rice susceptibility to bacterial blight. When the IE383 transgenic plants were treated with GA and DEX simultaneously, and susceptibility to bacterial blight was further enhanced. However, the DEX-treated IE383 transgenic line showed equal length of disease lesions on the *Xoo*-inoculated leaves to the wild-type plants after PAC treatment. The results indicate that PAC treatment overrides the immunosuppressive effect of SCRE9 and that SCRE9 and GAs might function synergistically in the same signaling pathway to inhibit rice immunity. Taken together, SCRE9 might cause rice plants more susceptibility to fungal and bacterial diseases by promoting GA signaling.

### OsSIP1 is targeted by SCRE9 and interacts with the transcription factors OsMADS63 and OsMADS68

To explore how SCRE9 regulates GA signaling, the SCRE9-interacting proteins were identified in the SCRE9-FLAG-expressing transgenic rice plants using immunoprecipitation-mass spectrometry. After immunoprecipitation with anti-FLAG M2 affinity beads, 252 candidate interactors of SCRE9 were identified in the anti-FLAG immunocomplex through mass spectrometry. Subsequently, *Oryza sativa* <u>S</u>CRE9-interacting <u>p</u>rotein 1 (OsSIP1, Os03g0100200) was identified as an interactor of SCRE9 from 45 tested proteins by yeast two-hybrid assays (Fig. 6A). This interaction was further confirmed by co-immunoprecipitation (co-IP) assays. After OsSIP1-HA was transiently co-expressed with SCRE9-FLAG in rice protoplasts, total proteins were extracted from the transfected protoplasts and were subjected to immunoprecipitation using anti-FLAG affinity beads. As detected by immunoblotting, OsSIP1-HA was detected in anti-SCRE9-FLAG immunocomplex (Fig. 6B).

**Figure 6.**
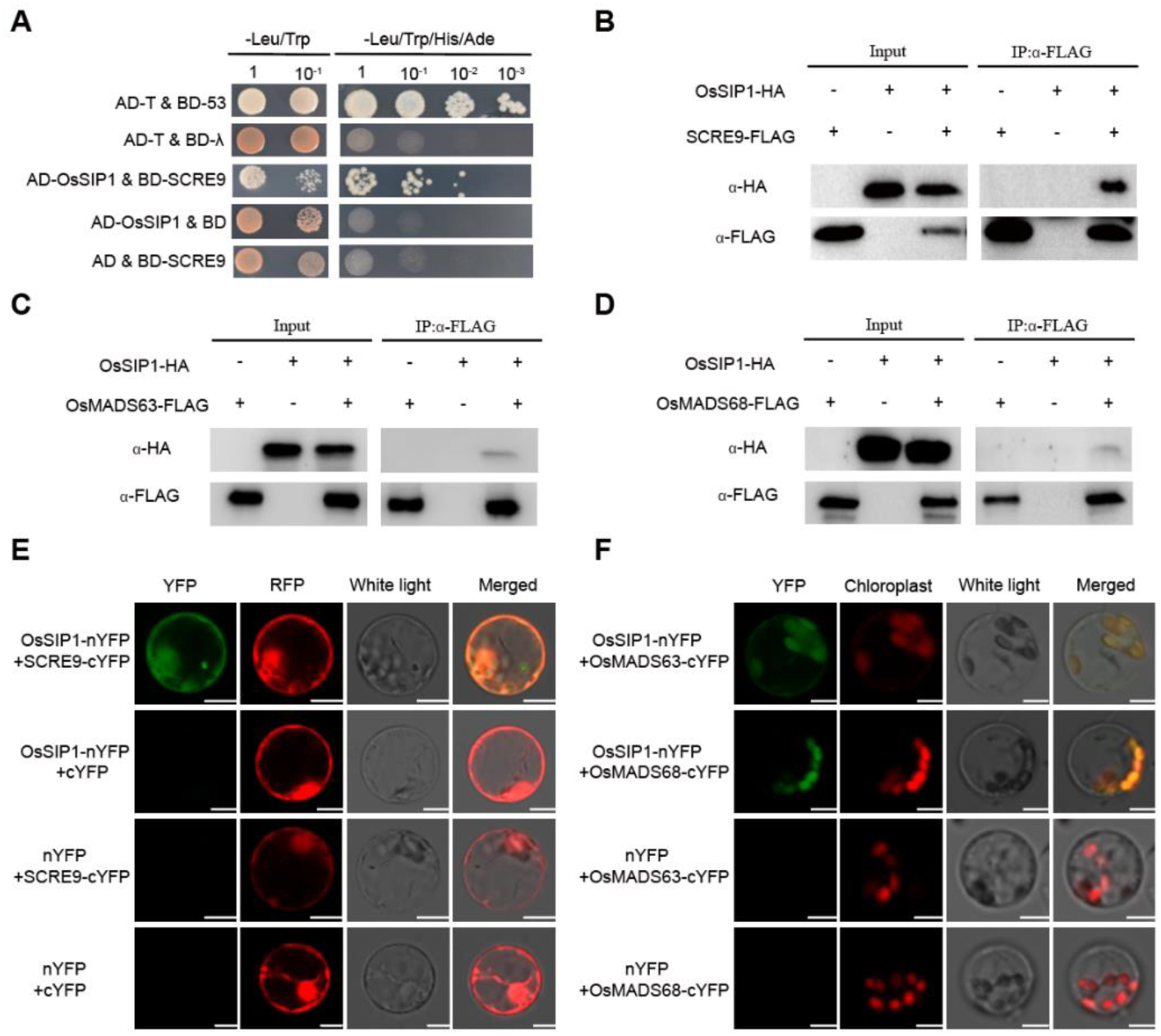
OsSIP1 is not only targeted by SCRE9 but also interacts with OsMADS63 and OsMADS68. A, Yeast two-hybrid assays to show the interaction of SCRE9 with OsSIP1. Positive interactions were evaluated on medium plates containing an SD base lacking Ade, His, Leu, and Trp (SD–LTAH). The pGADT7-T plasmid was transformed with pGBKT7-53 and pGBKT7-λ into yeast cells as positive and negative controls, respectively. BD, pGBKT7; AD, pGADT7. B-D, Co-IP assays to detect the specific interaction of OsSIP1 with SCRE9 (B), OsMADS63 (C), and OsMADS68 (D) in rice protoplasts. OsSIP1-HA was individually expressed or was co-expressed with SCRE9-FLAG, OsMADS63-FLAG, or OsMADS68-FLAG in rice protoplasts. Total protein extracts from transfected protoplasts were incubated with anti-FLAG M2 affinity beads. The input proteins and immunoprecipitates were detected with anti-FLAG and anti-HA antibodies. E-F, BiFC assays to detect the specific interaction of OsSIP1 with SCRE9 (E), OsMADS63 and OsMADS68 (F) in rice protoplasts. OsSIP1-nYFP was co-expressed with SCRE9-cYFP, OsMADS63-cYFP, or OsMADS68- cYFP in rice protoplasts. YFP signals were observed at 16 h after co-expression. As negative controls, OsSIP1-nYFP and cYFP were co-expressed while nYFP was co-expressed with SCRE9-cYFP, OsMADS63-cYFP, and OsMADS68-cYFP. YFP panels, yellow fluorescence; RFP panels, red fluorescence from RFP as a cytoplasmic and nuclear localization marker. Chloroplast panels, auto-fluorescence in the chloroplasts. Scale bar, 10 µm.

The transcription factors OsMADS63 and OsMADS68 function synergistically in flower development (Liu et al. 2013) and likely regulate GA biosynthetic and metabolic pathways in the GO biological process (https://geneontology.org/). Therefore, we tested whether OsSIP1 interacts with these MADS transcription factors. OsMADS63-FLAG or OsMADS68-FLAG was individually expressed or co-expressed with OsSIP1-HA in rice protoplasts. Protein extracts from the transfected protoplasts were subjected to immunoprecipitation with anti-FLAG affinity beads. As detected by immunoblotting, OsSIP1-HA was co-immunoprecipitated with MADS63-FLAG and MADS68-FLAG (Fig. 6, C and D). Next, OsSIP1-nYFP and SCRE9-cYFP were co-expressed in rice protoplasts for bimolecular fluorescence complementation (BiFC) assays. YFP fluorescence was observed in the nucleus and cytoplasm of rice protoplasts (Fig. 6E), indicating that SCRE9 interacts with OsSIP1 in the cytoplasm and nucleus of plant cells. In addition, the interactions of OsSIP1 with OsMADS63 and OsMADS68 were verified by BiFC. Interestingly, the results showed that OsSIP1 interacted with OsMADS63 and OsMADS68 in the chloroplasts (Fig. 6F). Collectively, SCRE9 targets OsSIP1 that interacts with MADS63 and MADS68.

### SCRE9 diminishes the inhibitory effect of OsSIP1 on *GA3ox1* expression

As a rate-limiting enzyme in the GA biosynthetic pathway, GA3ox1 plays a crucial role in GA synthesis. Two CArG-box elements in the promoter region of *GA3ox1* that might be bound by MADS transcription factors were predicted via PlantPan 4.0 (https://plantpan.itps.ncku.edu.tw/plantpan4/) (Supplementary Table S2). To determine whether OsMADS63 or OsMADS68 directly binds to the CArG-boxes, an electrophoretic mobility shift assay (EMSA) was performed. The biotin-labeled probes were incubated with in vitro-purified GST, GST-OsMADS63, and GST-OsMADS68 fusion protein. Shifted bands were observed when the labeled DNA probes were mixed with GST-OsMADS63 and GST-OsMADS68, indicating that these transcription factors bind to the CArG-boxes. By contrast, no shifted band was detected when GST or mutant probes were used. Shifted bands faded with increasing concentrations of un-labeled probes added, indicating that OsMADS63 and OsMADS68 specifically bind to the tested CArG-boxes in vitro (Fig. 7, A and B).

**Figure 7.**
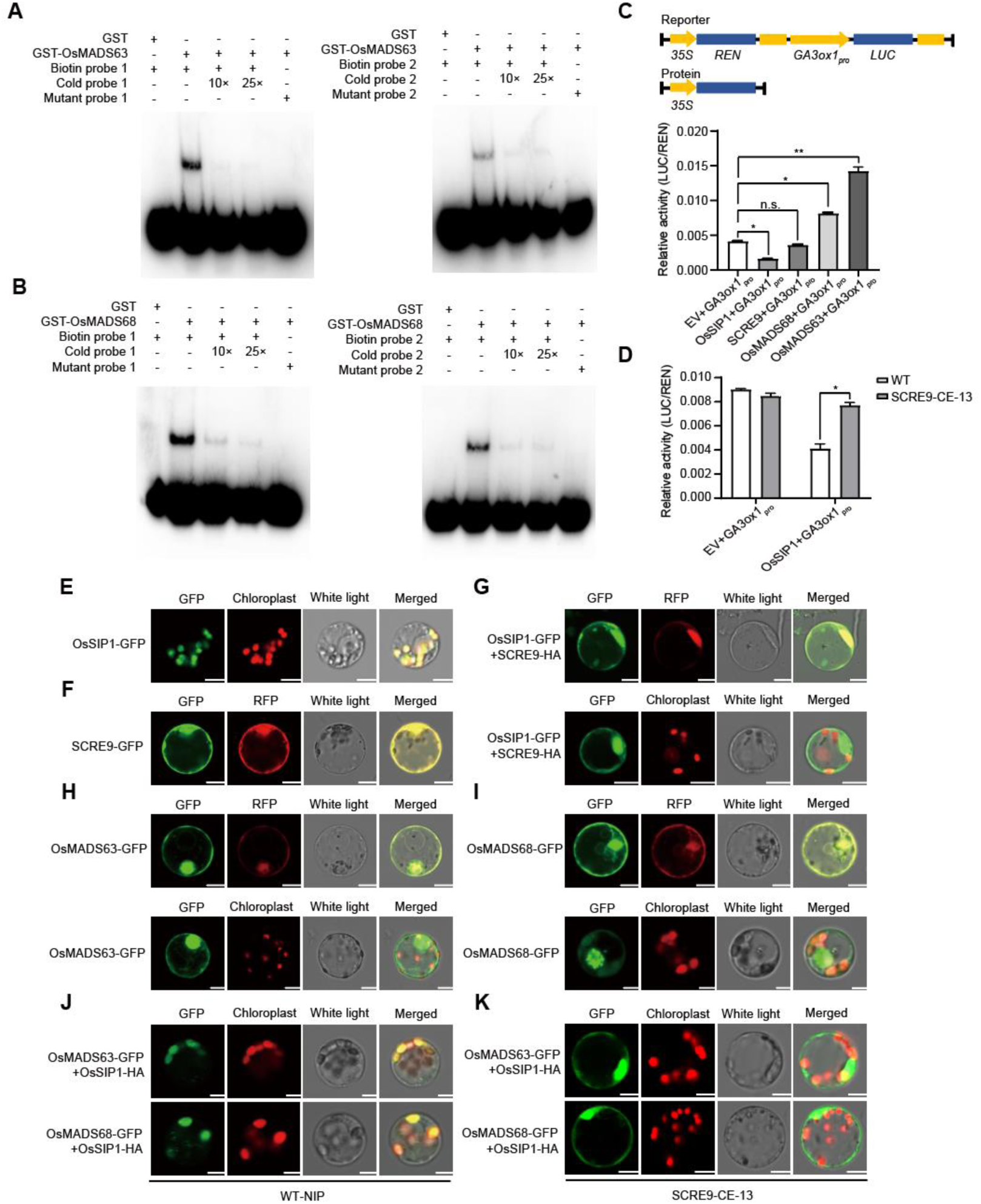
SCRE9 attenuates OsSIP1-mediated inhibition of *GA3ox1* expression through releasing OsSIP1-mediated confinement of OsMADS63/68 in the chloroplasts. A-B, EMSA to confirm the binding of OsMADS63 (A) or OsMADS68 (B) to the CArG-boxes. Purified GST-OsMADS63 or GST-OsMADS68 was incubated with biotin-labeled probes. The biotin-labeled probe (0.2 μM) was used, and unlabeled probes at different concentrations as indicated was added as competitors. GST and mutant probes were used as negative controls. C, Dual-LUC assays to show the LUC/REN activity when the *ProOsGA3ox1:LUC* construct was co-transfected with different gene constructs in rice protoplasts. Upper diagram shows the features of the constructs used in dual LUC assays. The LUC/REN activity was measured after *LUC* and different gene constructs were co-transfected into rice protoplasts. Asterisks indicate significant differences in the LUC/REN activity in rice protoplasts (One-way ANOVA least significant difference test; *, *P*<0.05; **, *P*<0.01; n.s., no significance). D, Dual LUC assays to show the LUC/REN activity when the *ProOsGA3ox1:LUC* construct was co-transfected with *OsSIP1* gene construct in the protoplasts of the wild-type NIP and SCRE9-expressing CE13 transgenic lines (*, *P*<0.05). Representative data from three independent assays are presented as mean ± SE (*n* = 4). E-F, Subcellular localization of OsSIP1-GFP (E) and SCRE9-GFP (F) in rice protoplasts. OsSIP1-GFP and SCRE9-GFP were individually expressed in rice protoplasts. G, Subcellular localization of OsSIP1-GFP when the protein was co-expressed with SCRE9-HA in rice protoplasts. H-I, Subcellular localization of OsMADS63-GFP and OsMADS68-GFP in rice protoplasts. OsMADS63-GFP and OsMADS68-GFP were individually expressed in rice protoplasts. J-K, Subcellular localization of OsMADS63-GFP and OsMADS68-GFP when these transcription factors were co-expressed with OsSIP1-HA in rice protoplasts isolated from the wild-type (J) and CE13 (K) transgenic lines. Fluorescence was observed via laser scanning confocal microscopy. GFP panels, green fluorescence; RFP panels, red fluorescence from RFP as a cytoplasmic and nucleus localization marker. Chloroplast panels, auto-fluorescence in the chloroplasts. Scale bar, 10 µm.

Next, we investigated the regulatory effects of SCRE9, OsSIP1, OsMADS63 and OsMADS68 on *GA3ox1* expression through dual luciferase (Dual-LUC) reporter assays in rice protoplasts. The firefly LUC reporter was expressed under the control of the *GA3ox1* promoter, while Renilla luciferase (REN) was expressed under the 35S promoter as an internal reference. The results showed that the relative reporter activity (LUC/REN) was significantly enhanced by co-expression of OsMADS63 and OsMADS68. In contrast, the LUC/REN activity was significantly inhibited by OsSIP1 co-expression (Fig. 7C). These data indicate that both MADS63 and MADS68 transcriptionally induce the expression of *GA3ox1*, whereas OsSIP1 suppresses the expression of *GA3ox1*. In dual LUC assays, the LUC/REN activity was not altered by SCRE9 expression, indicating that the effector had no direct effect on *GA3ox1* expression. To explore whether the SCRE9-OsSIP1 interaction has any effect on the expression of *GA3ox1*, the dual LUC assay was conducted using the protoplasts isolated from the wild-type and SCRE9-expressing transgenic lines. Interestingly, the LUC/REN activity was inhibited by transiently expressed OsSIP1 in the wild-type protoplasts, whereas transient expression of OsSIP1 did not inhibit LUC/REN activity in the SCRE9-expressing rice protoplasts (Fig. 7D). These data indicate that ectopic expression of SCRE9 diminishes the inhibitory effect of OsSIP1 on *GA3ox1* expression.

### SCRE9 releases OsSIP1-mediated restriction of OsMADS63/68 in the chloroplasts

To explore how SCRE9 interferes with OsSIP1 function in inhibiting *GA3ox1* expression, subcellular localization of these proteins was investigated through transient gene expression in rice protoplasts and in *Nicotiana benthamiana* leaves. Transiently expressed OsSIP1-GFP was exclusively localized in the chloroplasts (Fig. 7E; Supplementary Fig. S5A), while SCRE9-GFP was mainly localized in the cytosol and nucleus, and was also partly present in the chloroplasts (Fig. 7F; Supplementary Fig. S5B). Interestingly, OsSIP1-GFP was relocated into the nucleus and cytoplasm when OsSIP1-GFP was co-expressed with SCRE9-HA in rice protoplasts (Fig. 7G; Supplementary Fig. S5C). In addition, we demonstrated that OsMADS63-GFP and OsMADS68-GFP were localized in the nucleus and cytoplasm when these proteins were individually expressed in rice protoplasts and *N. benthamiana* leaves (Fig. 7, H and I; Supplementary Fig. S5D). However, both OsMADS63 and OsMADS68 were partially recruited into the chloroplasts when OsSIP1-HA was co-expressed with OsMADS63 and OsMADS68 in rice and *N. benthamiana* (Fig. 7J; Supplementary Fig. S5E). Notably, chloroplast localization of OsMADS63 and OsMADS68 recruited by OsSIP1 was greatly reduced in the rice protoplasts of the CE13 transgenic line. OsMADS63 and OsMADS68 were mainly relocated back to the nucleus and cytoplasm (Fig. 7K). These results indicate that OsSIP1 inhibits the transcriptional activation of *GA3ox1* by restricting the transcription factors OsMADS63 and OsMADS68 in the chloroplasts, while the effector SCRE9 interacts with OsSIP1 and releases OsSIP1-mediated restriction of OsMADS63 and OsMADS68 in the chloroplasts.

## Discussion

Cell walls in stamen filaments might function as a major physical barrier for *U. virens* infection because the pathogen does not form specialized feeding structures (e.g. haustoria) and appressoria, and is deficient in the ability to degrade cell wall (Fan et al. 2016;Tang et al. 2013; Zhang et al. 2014). In this study, we identify a crucial *U. virens* virulence effector SCRE9 that promotes GA biosynthesis and induces expression of expansins and β-glucosidases in rice panicles, which might contribute to loosen cell walls and break physical barriers for *U. virens* infection.

SCRE9 has characteristics of an effector protein. First, *SCRE9* is transcriptionally induced during *U. virens* infection. Second, we confirm the secretory function of SCRE9 signal peptide (Fig. 1A). Many effectors in filamentous fungi are secreted into the extracellular spaces of host cells under the guidance of signal peptides, and are then translocated into plant cells (Dou and Zhou 2012). Multiple effector proteins in *U. virens* have been demonstrated to be translocated into host cells through the BICs when the effectors were heterologously expressed in *M. oryzae* (Qiu et al. 2022; Zheng et al. 2022). Here, ectopically expressed SCRE9-GFP in *M. oryzae* has been also demonstrated to be accumulated in the BICs during infection, indicating that SCRE9-GFP is secreted and translocated into host cells via the BICs (Supplementary Fig. S1C). Furthermore, *SCRE9* knockout causes *U. virens* less virulent (Fig. 1, B and C). Expression of SCRE9 in transgenic rice lines causes an evident suppression of PAMP-triggered immune responses and resistance to *U. virens* and *X. oryzae* pv. *oryzae* (Fig. 2; Fig. 5). Taken together, these results demonstrate that SCRE9 is an essential virulence factor in *U. virens*.

Next, we elucidate molecular mechanisms how SCRE9 inhibits rice immunity through comparison of gene expression profilings between the wild-type and SCRE9-expressing transgenic lines. GA synthesis-related genes are enriched in the DEGs that are up-regulated by SCRE9 (Fig. 4, B and C). Consistently, these transgenic plants exhibit robust growth with elevated plant height. More convincingly, endogenous gibberellins, particularly active gibberellins, are elevated in the SCRE9-expressing panicles (Fig. 4, D and E). Therefore, we hypothesize that SCRE9 inhibits floret immunity through regulating GA biosynthesis in rice. GA, as a growth stimulating hormone, has recently identified as a negative regulator of plant immune signaling (De Vleesschauwer et al. 2013; De Vleesschauwer et al. 2016; Shahnejat-Bushehri et al. 2016; Gao et al. 2017). Besides antagonizing with SA- and JA-mediated defense signaling (De Vleesschauwer et al. 2013; Van der Does et al. 2013; Zhang et al. 2023), GA induces the expression of expansin genes (Lee and Kende 2001; Azeez et al. 2010; Sun et al. 2011; Li et al. 2019). Expansins can lead to microfibril dispersion and cell wall loosening, which is conducive to pathogen infection (Cosgrove 2000; Cosgrove 2024). Silencing of α-expansin 4 gene (*EXPA4*) enhances *N. benthamiana* resistance to tobacco mosaic virus, whilst *EXPA4* overexpression promotes virus reproduction on tobacco (Chen et al. 2018). Tomato fruit-specific expansin LeExp1 promotes susceptibility to the fungal pathogen *Botrytis cinerea* (Cantu et al. 2009). *X. oryzae* pv. *oryzae* infection induces expression of expansin genes in rice, thus causing cell wall loosening and facilitating the invasion of pathogenic bacteria (Ding et al. 2008). In this study, a number of expansin, pollen allergen and β-glucosidase genes have been identified to be induced in the SCRE9-expressing young panicles (Supplementary Table S1). Elevated accumulation of active GAs and induced expression of expansins and β-glucosidases could loosen cell walls in floret filament, which helps break the physical barrier for infection of *U. virens*. The statement is further substantiated by other experimental data. Exogenous application of GA causes rice plants more susceptibility to various pathogens. By contrast, treatment of the GA synthesis inhibitor PAC makes rice plants more resistant. Meanwhile, PAC treatment completely eliminates the enhanced susceptibility caused by SCRE9, indicating that SCRE9-mediated immunosuppression requires the action of GA (Fig. 5). Altogether, SCRE9 inhibits floral immunity and promotes *U. virens* infection by inducing GA synthesis and expression of expansins and pollen allergens in rice.

Furthermore, the regulatory mechanism of SCRE9 on GA biosynthesis was revealed by subcellular localization observation and protein-protein interaction analyses. The SCRE9-OsSIP1 interaction was identified and confirmed by various approaches including IP-MS, yeast two-hybridization and co-IP assays (Fig. 6, A and B). Interestingly, OsSIP1 interacts with the transcription factors OsMADS63 and OsMADS68 (Fig. 6, C and D), which function synergistically in flower development (Liu et al. 2013). We further show that OsMADS68 and OsMADS63 transcriptionally induce the expression of *GA3ox1* encoding a rate-limiting enzyme in GA biosynthesis (Fig. 7). Dual LUC assays show that OsSIP1, but not SCRE9, suppresses the expression of *GA3ox1*, whereas SCRE9 expression in the transgenic lines eliminates the inhibitory effect of OsSIP1 on *GA3ox1* expression (Fig. 7).

Some chloroplastic proteins have also been reported to regulate GA content. CND41 is a nucleoid DNA binding protein in the chloroplast and acts as a negative regulator of active gibberellin levels (Nakano et al. 2003). Overexpression of *GmDXRs* (plant 1-deoxy-D-xylulose 5-phosphate reductoisomerase) induces an increase in the contents of various isoprenes (chlorophyll, carotenoid and gibberellin), and regulates isoprene biosynthesis (Zhang et al, 2012). Overexpression of *OsAL13* in the chloroplast leads to increased GA3 content in the shoot (Guo et al, 2024). However, it remains to be identified how these chloroplastic proteins are involved in regulating gibberellin biosynthesis.

Subcellular localization observation provides an explanation how SCRE9, OsSIP1, OsMADS68 and OsMADS63 regulate GA biosynthesis in rice. As transcription factors, OsMADS68 and OsMADS63 are mainly localized into the nuclei. In contrast, OsSIP1 is localized in the chloroplasts although the protein carries no chloroplast localization signal (Fig. 7, E, H, and I). Notably, the majority of OsMADS63 and OsMADS68 are translocated into the chloroplasts when these transcription factors are co-expressed with OsSIP1 in rice protoplasts (Fig. 7J). Based on these findings, we hypothesize that OsSIP1 recruits OsMADS63 and OsMADS68 into the chloroplasts and reduces transcription activation of GA biosynthetic genes, such as *OsGA3ox1*, and GA biosynthesis (Fig. 8). Furthermore, we show that SCRE9 co-expression causes OsSIP1 translocation from the chloroplasts into the nucleus through unknown mechanism, perhaps by physical interaction (Fig. 7; Supplementary Fig. S5). More interestingly, OsMADS63 and OsMADS68 are mainly localized into the nuclei of the SCRE9-expressing transgenic rice protoplasts even when these proteins are co-expressed with OsSIP1 (Fig. 7K). Therefore, during *U. virens* infection, the secreted SCRE9 interacts with OsSIP1, thereby releasing OsSIP1-mediated confinement of OsMADS63 and OsMADS68 in the chloroplasts and promoting the transcriptional activation of GA biosynthetic genes such as *GA3ox1* in the nuclei (Fig. 8).

**Figure 8.**
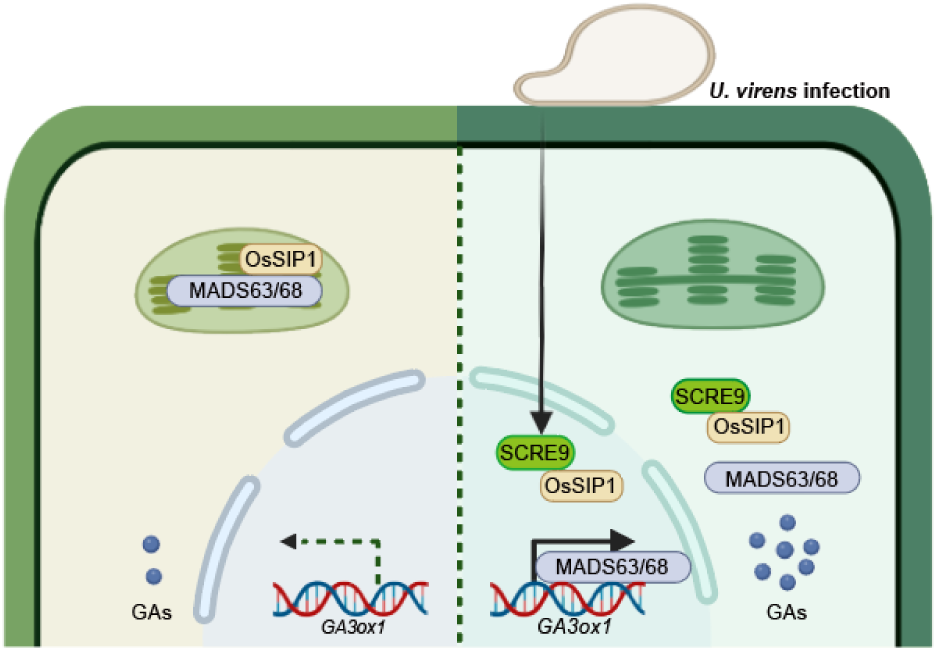
A working model of SCRE9 promoting GA biosynthesis and suppressing floret immunity in rice. OsMADS63 and OsMADS68 promote the expression of *GA3ox1* encoding a key enzyme in GA synthesis. OsSIP1 is localized in the chloroplasts and simultaneously recruits OsMADS63 and OsMADS68 into the chloroplasts, thus reducing transcriptional activation of *GA3ox1*. During *U. virens* infection, the effector SCRE9 is secreted and translocated into rice nucleus and cytoplasm. SCRE9 recruits OsSIP1 into the nucleus and cytoplasm, thereby releasing OsMADS63 and OsMADS68 from the chloroplasts into cell nuclei. These transcription factors subsequently enhance the expression of GA biosynthetic genes including *GA3ox1*, thus promoting the synthesis of GAs and inhibiting floret immunity.

Plant hormones, such as SA, JA and ethylene, are extensively involved in regulating immune responses. A number of fungal effectors have been identified to target SA and JA signaling pathways to promote fungal infection. To our knowledge, this is the first report that a fungal effector facilitates fungal infection by promoting GA signaling pathway in plants. SCRE9 promotes the expression of GA synthesis genes in rice and increases the content of endogenous GAs. GAs induce the expression of expansin genes, thus loosening cell walls in floret filaments and facilitating *U. virens* infection. This study provides a theoretical basis for the understanding of pathogenic mechanism of *U. virens* specifically inhibiting rice floret immunity. Because *U. virens* is defective in cell wall-degrading ability, the action of SCRE9 represents an important type of invasion strategy for this biotrophic fungi.

## Materials and methods

### Microbial Strains and Plant Materials

The *U. virens* isolates P1FZ and JS60-2 were cultured in potato sucrose agar (PSA, boiled extracts from 200 g of fresh potato, 20 g of sucrose, and 14 g agar per liter) plates at 28°C. P1FZ was used to generate the *scre9* knockout mutants. *M. oryzae* Guy11 and *X. oryzae* pv. *oryzae* PXO99 were cultured in oatmeal–tomato paste medium (boiled extracts from 30 g of oats, 150 mL tomato juice, and 14 g agar per liter) and nutrient broth (NB) at 28°C, respectively. *Escherichia coli* and *Agrobacterium tumefaciens* strains were cultured on Luria–Bertani (LB) medium at 37°C and 28°C, respectively. The rice cultivar Nipponbare (NIP) was used as the wild type in this study. NIP, Jinong853 (JN853) and the derivative transgenic rice plants were grown in the greenhouse for pathogen inoculation assays.

### RNA Isolation and Quantitative RT-PCR

Total RNAs were isolated from rice seedlings, rice panicles and *U. virens* mycelia using an Ultrapure RNA kit (CWBio) following the manufacturer’s instructions. RNAs were quantified via NanoDrop One (Thermo Fisher Scientific). Complementary DNA was synthesized via the M-MLV reverse transcriptase system (TaKaRa) using total RNAs as template and random oligonucleotides as primers. Quantitative RT-PCR was performed with a LightCycler^R^ 96 Instrument (Roche) using gene-specific primers.

### Yeast Secretion Assay

The yeast secretion assay was performed as described previously (Zou et al. 2023). The predicted signal peptide (SP)-coding sequence of *SCRE9* was subcloned into the vector pSUC2. The resultant pSUC2-*SCRE9^SP^* plasmid was transformed into the yeast strain YTK12. The enzymatic activity of secreted invertase was detected by the reduction of TTC (2,3,5-triphenyltetrazolium chloride) to insoluble red TPF (1,3,5-triphenylformazan).

### Effector Translocation Assay

Effector translocation was detected as described previously (Qiu et al. 2022; Zheng et al. 2022). The coding sequence of *SCRE9* was amplified and subcloned into the modified pYF11-*RP27Pro:GFP* vector via homologous recombination. The plasmid was transformed into *M. oryzae* Guy11 via PEG-mediated protoplast transformation. The transformants with strong green fluorescence were selected for inoculation assays. Conidial suspension (1×10^5^ spores/mL) was prepared and injection-inoculated into rice leaf sheaths. The inoculated leaf sheaths were incubated in a growth chamber in darkness at 28°C for 30-36 h. Green fluorescence in the BICs was observed under confocal microscopy (Leica STELLARIS 5, Germany).

### Southern Blot Analysis

Southern blot analysis was performed as described previously (Fang et al. 2019; Zou et al. 2023). Briefly, genomic DNA was isolated from the wild-type strain and *Δscre9* knockout candidates using CTAB extraction buffer (Sangon Biotech) and was then digested with *Sac* I and *Sph* I overnight. Probe labeling and signal detection were performed using a DIG High Prime DNA Labeling Kit and a Detection Starter Kit (Roche), respectively, according to the manufacturer’s instructions.

### ROS burst assay

PAMP-induced ROS burst was detected as described previously (Fang et al. 2019; Zhang et al. 2020). The 6-week-old wild-type and transgenic plants were sprayed with DEX (10 μM) or mock solution. Leaf discs were collected from the treated plants and were then incubated in sterile water overnight. Leaf discs were immersed in 100 μM of D-luciferin potassium salt substrate (Gold Biotechnology) supplemented with flg22 (1 μM), chitin (10 µg/mL), or sterile deionized water. ROS generation was monitored by a GloMax 20/20 luminometer (Promega) immediately for 25 min.

### RNA sequencing

The wild-type and transgenic rice panicles were injected with DEX solution (10 μM) at 5-7 days before heading, and were then injection-inoculated at 16 h after DEX treatment. RNA-seq was performed using a second-generation high-throughput sequencing Illumina platform by Gene Denovo Biotechnology Co. (Guangzhou, Guangdong). The genes showing |Log2(Fold Change)| ≥ 1 and an adjusted *P*-value < 0.05 were considered as differentially expressed genes (DEGs). A Venn diagram was plotted and PCA was performed using the online Omicsmart tools at https://www.omicsmart.com/. All DEGs were mapped to GO terms in the Gene Ontology database to determine the main biological functions (http://www.geneontology.org/). KEGG pathway enrichment analysis was performed to identify the metabolic pathways and signal transduction pathways significantly enriched for DEGs.

### Determination of GA content

Endogenous GAs in the DEX-treated wild-type and transgenic rice panicles were determined by UHPLC-MS/MS analysis (Thermo Scientific Ultimate 3000 UHPLC coupled with TSQ Quantiva) at Wuhan Greensword Creation Technology Co. Ltd. (Wuhan, Hubei, China).

### *Ustilaginoidea virens* inoculation assay

*U. virens* inoculation assay was performed as described previously (Qiu et al. 2022; Zheng et al. 2022). Briefly, *U. virens* was cultured on PSA plates at 28°C for 10 d, and was then transferred into PSB liquid medium. After culturing for one week with shaking at 150 rpm at 28°C, mycelial masses were smashed with a blender. The mixture of hyphae and spores were diluted to 1×10^6^ spores/mL using PSB medium and were then injected into rice panicles at 5 – 7 days before heading. At least 10 panicles were inoculated for each strain. False smut balls formed on rice panicles were counted at 4 weeks after inoculation.

### Bacterial inoculation assay

Six-week-old rice plants were inoculated with *X. oryzae* pv. *oryzae* PXO99A using the leaf-clipping method (Zheng et al. 2022; Zou et al. 2023). The leaf tip was cut using a pair of scissors soaked in cell suspension of *X. oryzae* pv. *oryzae* (OD600 = 0.8). The length of disease lesions on the inoculated leaves was measured at 14 d post inoculation.

### Yeast two-hybrid assay

The coding sequences of *SCRE9* and *OsSIP1* were subcloned into pGBKT7 (BD) and pGADT7 (AD), respectively. The pair of AD and BD constructs for each potential interaction were co-transformed into the yeast Gold strain using a Frozen-EZ Yeast Transformation II Kit (ZYMO RESEARCH). The transformants were screened on synthetic defined (SD) dropout (without Leu and Trp, SD/–Leu–Trp) medium plates. The interaction was determined after the transformants were cultured on SD base quadruple dropout medium (SD/–His–Ade–Leu–Trp) plates. The plasmid AD-T was co-transformed with BD-53 and BD-λ into yeast cells as positive and negative controls, respectively.

### Co-IP assay

The coding sequences of *SCRE9* and *OsSIP1* were subcloned into pUC19-35S-FLAG and pUC19-35S-HA, respectively. Rice protoplasts were isolated from the seedlings as described previously (Yang et al. 2022; Zheng et al. 2022). The constructed vectors were co-transfected into rice protoplasts (2 × 10^6^ cells/mL). Total proteins extracted from transfected protoplasts were incubated with anti-FLAG M2 affinity beads (Sigma-Aldrich, A2220) at 4°C for 4 h. The beads were rinsed thoroughly with IP buffer and the immunoprecipitants were eluted using 1 × SDS–polyacrylamide gel electrophoresis (PAGE) loading buffer. The samples were subjected to immunoblot analyses using an HRP-conjugated anti-HA antibody (Roche, 11667475001, 1: 2,000 dilution) and an anti-FLAG antibody (Sigma-Aldrich, F1804, 1: 5,000 dilution).

### BiFC assay

The coding sequences of *OsSIP1* and *SCRE9*, *OsMADS63* or *OsMADS68* were subcloned into the vectors pRTV-nYFP and pRTV-cYFP, respectively (Wei et al. 2020). The *OsSIP1-nYFP* construct was co-transfected with *SCRE9-cYFP*, *OsMADS63-cYFP* or *OsMADS68-cYFP* into rice protoplasts. YFP fluorescence signal was observed in rice protoplasts at 16 h after transfection using confocal microscopy with laser excitation at a 514-nm wavelength and emissions ranged from 530 to 580 nm (Leica STELLARIS 5, Germany).

### Dual-LUC Reporter Assay

The dual-LUC reporter assay was performed as described (Qiu et al. 2022). Briefly, the promoter of *GA3ox1* was subcloned into pGreenII 0800-*LUC* to drive expression of the firefly luciferase gene (*LUC*). The expression of the Renilla luciferase gene (*REN*) under the control of 35S promoter was used as a reference. The *SCRE9*, *OsSIP1*, *MADS63* and *MADS68* genes were subcloned into pGreenII 62-SK. The constructed vectors were co-transfected into rice protoplasts isolated from the wild-type and transgenic rice seedlings. Luminescence was detected in rice protoplasts using a DualLumi^TM^ II luciferase Assay Kit (YEASEN, Shanghai, China) according to the manufacturer’s instructions. The relative luciferase activity was indicated by the ratio of LUC/REN.

### Subcellular Localization

The coding sequences of *SCRE9*, *OsSIP1*, *MADS63* and *MADS68* were amplified and subcloned into pRTV-GFP. The constructed vectors were transfected into rice protoplasts isolated from the wild-type and transgenic rice seedlings. Red and green fluorescence was observed in rice protoplasts using a Leica confocal microscope (Leica STELLARIS 5, Germany).

### Statistical Analysis

Significance analyses in multiple comparisons were performed with one-way ANOVA followed by Duncan’s multiple range test using SPSS software. Pairwise comparisons were evaluated through Student’s *t* test.

### Accession numbers

Sequence data of the rice genes studied in this article can be found in the rice genome annotation project database under the following accession numbers: *OsSIP1* (Os03g0100200); *OsMADS63* (Os06g0223300); *OsMADS68* (Os11g0658700); *GA3ox1* (Os05g0178100).

## Author contributions

W.S., D.L., and S.Y. conceived this study. S.Y., S.Z., X.L., X.Z., J.W., G.D., S.Q., D.Z., N.N., Q.Y., C.J., P.Z. performed assays and analyzed data. W.S., D.L., and S.Y. wrote and edited the article. All authors discussed the results, reviewed and approved the article.

## Supplemental data

The following materials are available in the online version of this article.

Supplemental Figure S1. SCRE9 encoding a putative secreted protein is transcriptionally induced during *U. virens* infection.

Supplemental Figure S2. The *SCRE9* knockout and complemented strains are confirmed by Southern and western blot analyses.

Supplemental Figure S3. DEX-induced expression of SCRE9 inhibits PAMP-triggered ROS burst.

Supplemental Figure S4. Constitutive expression of SCRE9 promotes growth and development in rice.

Supplemental Figure S5. OsSIP1 recruits MADS63 and MADS68 into chloroplasts and SCRE9 recruits OsSIP1 into the nucleus.

Supplementary Table S1. The up-regulated genes encoding expansins, pollen allergens and β-glucosidases in the IE383 transgenic panicle.

Supplementary Table S2. Probes used in EMSA.

Supplementary Table S3. Primers used in this study.

## Acknowledgments

We thank Zhengguang Zhang at Nanjing Agricultural University for the yeast strain XK-125, *M. oryzae* Guy11, and the pYF11 vector; Jinrong Xu at Purdue University for the pmCAS9-tRp-gRNA vector; Zhaoxi Luo at Huazhong Agricultural University for the JS60-2 isolate; Fangjun Li at China Agricultural University for pGreenII 0800-LUC vector and Wuhan Greensword Creation Technology Co. Ltd for providing GA content determination.

## Funding information

This work is supported by the National Natural Science Foundation of China (grants 32293241 and 32430090), the China Agricultural Research System (CARS01), Hainan Seed Industry Laboratory and China National Seed Group (project of B23YQ1514/B23CQ15EP), the Science and Technology Development Project of Jilin Province (20240303006NC and 20240304122SF).

## Conflict of interest statement

The authors declare no conflict of interests.

## Data availability

The data underlying this article are available in the article and/or the online supplementary materials.

---

**Figure S1.**
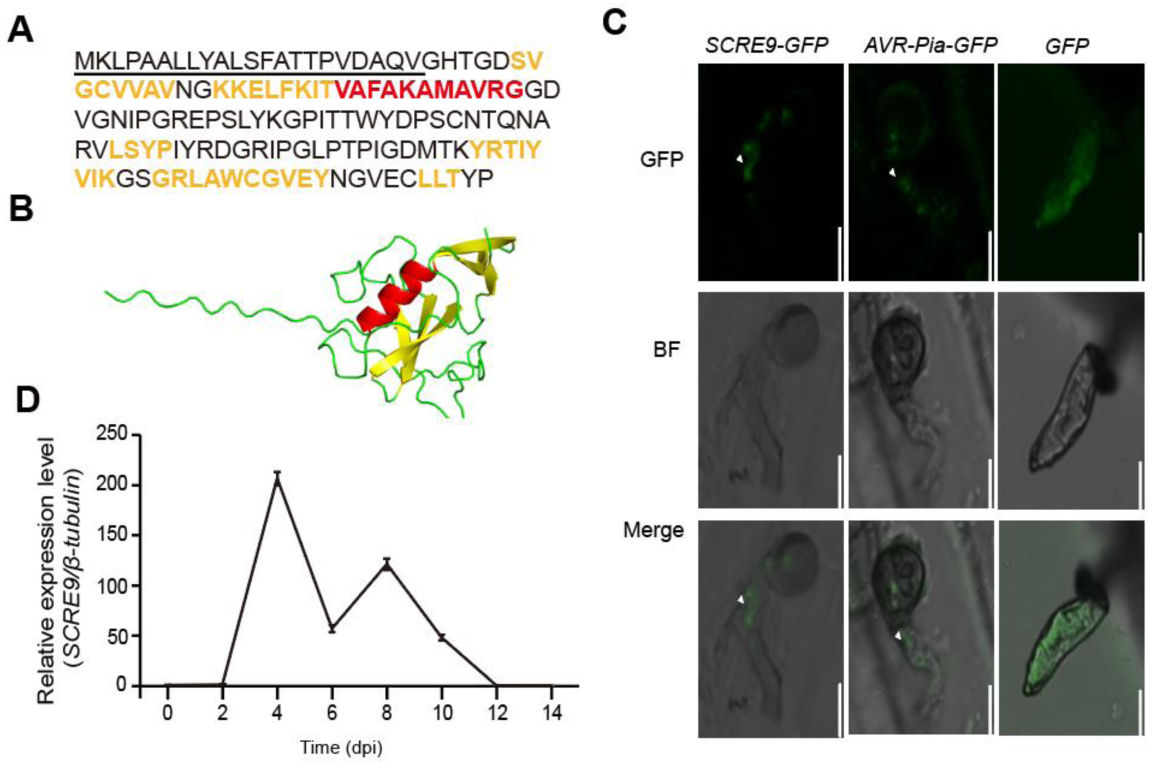
*SCRE9* encoding an effector is transcriptionally induced during *U. virens* infection. A-B, The predicted protein sequences and 3D structure of SCRE9 with typical structural features. The putative N-terminal signal peptide is underlined. The 3D structure of SCRE9 was predicted by AlphaFold2. The predicted sequences forming α-helixes (red color) and β-strands (orange color) are highlighted in red and orange, respectively. C, GFP fluorescence resulting from ectopic expression of SCRE9-GFP in *M. oryzae* was observed in the BICs at 30–36 h after *M. oryzae* inoculation. BICs are indicated by white triangles. GFP, green fluorescent protein; BF, bright field; Merge, the overlay of GFP and BF images. D, The expression pattern of *SCRE9* during *U. virens* infection. The expression of *SCRE9* was detected in the inoculated panicles via quantitative RT-PCR at the indicated time points after the susceptible rice variety JN853 was inoculated with *U. virens*. dpi, days post inoculation. The β-tubulin gene was used as an internal reference. The representative data from three independent experiments are presented as mean ± SE (*n* = 3).

**Figure S2.**
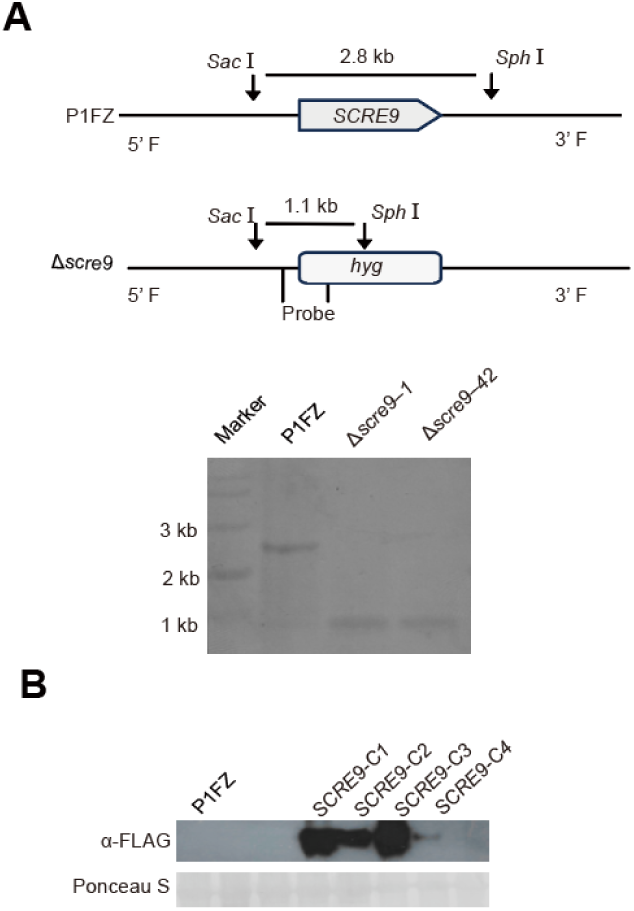
Southern and western blot analyses to confirm the *SCRE9* knockout and complemented strains, respectively. A, Southern blot analysis to confirm the Δ*scre9* deletion mutants. Upper panel, a schematic diagram to show restriction enzyme digestion and probe design for Southern blot analysis. Lower panel, the deletion mutants Δ*scre9-*1 and Δ*scre9-*42 were confirmed by Southern blot analysis. P1FZ, the wild-type strain. B, Immunoblot analysis to confirm the Δ*scre9-C* complemented strains. The Δ*scre9*-*C* complemented strains were screened by immunoblotting probed with an anti-FLAG antibody.

**Figure S3.**
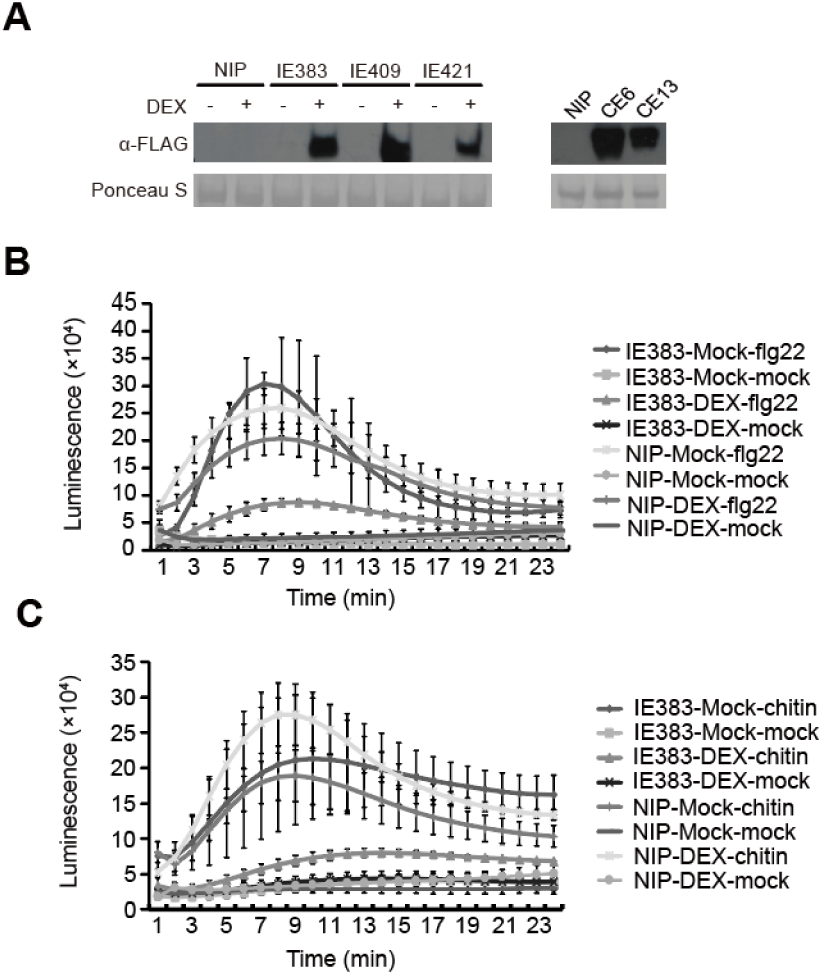
DEX-induced expression of SCRE9 inhibits PAMP-triggered ROS burst. A, The expression levels of SCRE9 in different transgenic rice lines as detected by immunoblotting. The IE383, IE409 and IE421 independent transgenic lines with DEX-inducible expression of SCRE9-FLAG were treated with DEX for 24 h. CE6 and CE13, the transgenic lines with constitutive expression of SCRE9. Immunoblotting was performed with an anti-FLAG antibody (α-FLAG). Nip, Nipponbare; DEX, dexamethasone; Mock, no DEX treatment. B-C, flg22- and chitin-triggered ROS burst in the wild-type and IE383 transgenic lines with or without DEX treatment. The wild-type and transgenic seedlings were treated with DEX (10 µM) or Mock solution for 24 h followed by flg22 (1 µM), chitin (10 µg/mL) or mock treatment. ROS burst was detected immediately after PAMP treatments for 24 min. The representative data from three independent experiments are presented as mean ± SE (*n* = 3).

**Figure S4.**
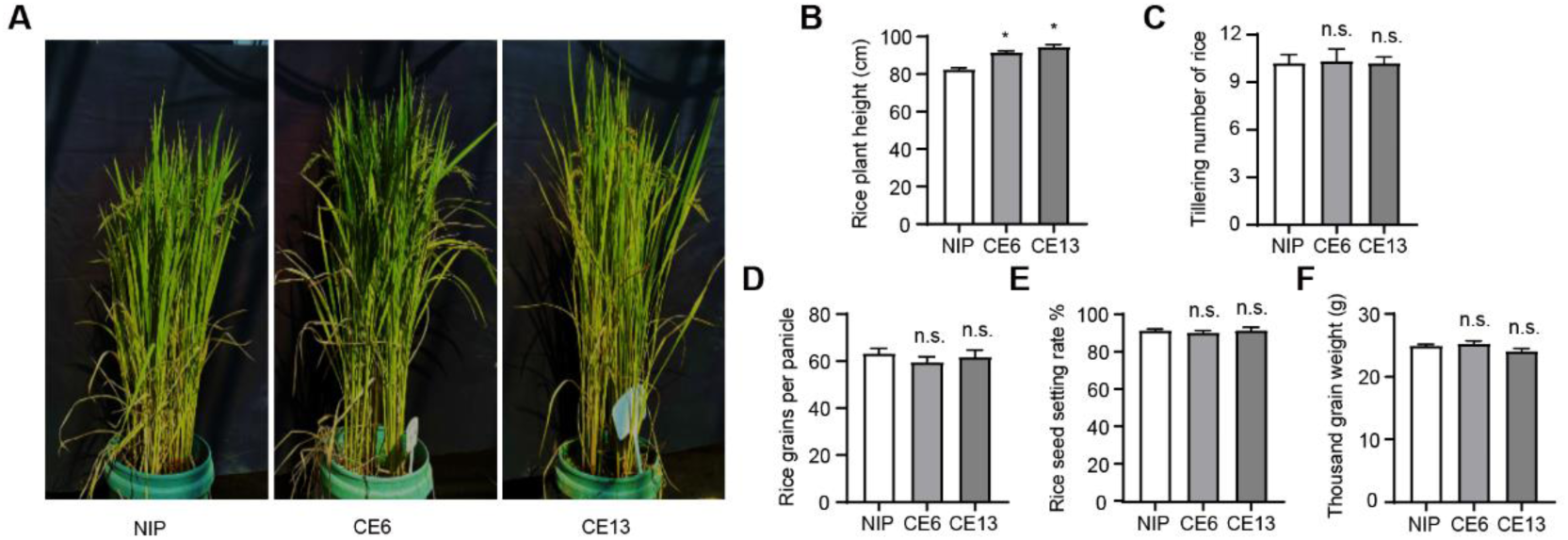
Constitutive expression of SCRE9 promotes rice growth. A, Growth phenotypes of the wild-type and SCRE9-expressing CE6 and CE13 transgenic rice plants. The images were taken in the greenhouse. B-F, The plant height (B), tillering number (C), grains per panicle (D), seed setting rate (E) and 1000-grain weight (F) of the wild-type and transgenic plants with constitutive expression of SCRE9. Representative data from three independent experiments are presented as mean ± SE (*n* = 15).

**Figure S5.**
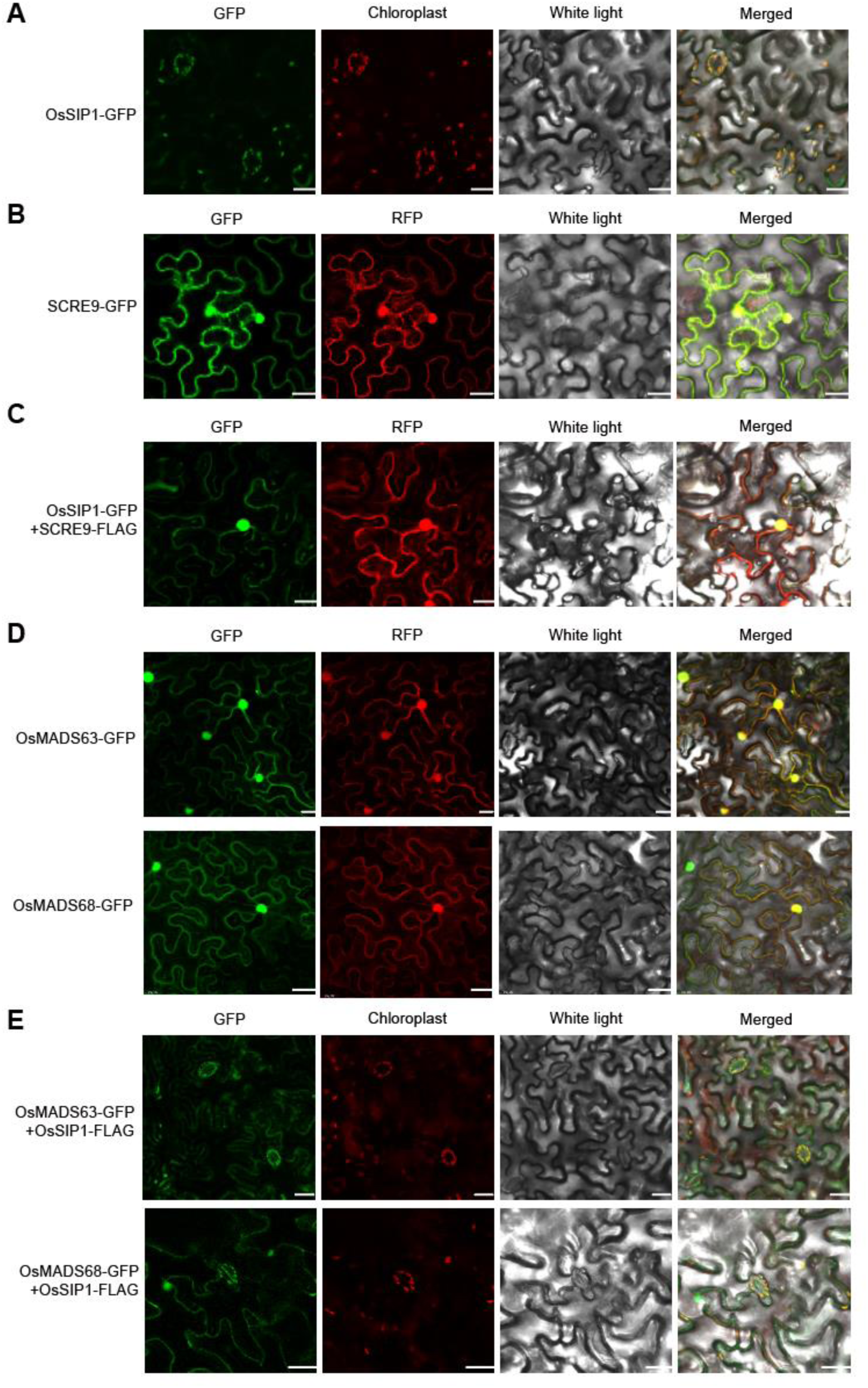
SCRE9 releases OsSIP1-mediated confinement of OsMADS63/68 in the chloroplasts of *Nicotiana benthamiana*. A, Subcellular localization of OsSIP1-GFP in *N. benthamiana* cells. OsSIP1-GFP was localized into the chloroplasts. B, Subcellular localization of SCRE9-GFP in *N. benthamiana*. SCRE9-GFP was localized into the nuclei and cytoplasm. C, Subcellular localization of OsSIP1-GFP after co-expression with SCRE9-FLAG in *N. benthamiana*. D, Subcellular localization of OsMADS63-GFP and OsMADS68-GFP in *N. benthamiana*. OsMADS63-GFP and OsMADS68-GFP were localized into nuclei and cytoplasm. E, Subcellular localization of OsMADS63-GFP and OsMADS68-GFP after co-expression with OsSIP1-FLAG in *N. benthamiana*. GFP panels: green fluorescence; Chloroplast panels: auto-fluorescence in the chloroplasts. RFP panels: red fluorescence as a cytoplasmic and nuclear localization marker. Scale bar, 10 µm.

**Supplementary Table S1.** The up-regulated genes encoding expansins, pollen allergens and β-glucosidases in the IE383 transgenic panicle.

| Gene ID | Gene description | Log<sub>2</sub>(fold change) | FDR |
| --- | --- | --- | --- |
| Os06g0662900 | pollen allergen Lol p 2-A | 6.087054782 | 0.0020297 |
| Os06g0662600 | pollen allergen Lol p 2-A | 5.318584652 | 6.81E-05 |
| Os06g0662850 | pollen allergen Phl p 2 | 4.784750303 | 6.81E-05 |
| Os06g0662500 | pollen allergen Phl p 2 | 4.75010878 | 0.026988205 |
| Os06g0655200 | pollen allergen Lol p 2-A | 4.21803076 | 0.026988205 |
| Os04g0329200 | pollen allergen Lol p 2-A | 3.911305228 | 0.009090438 |
| Os04g0317500 | pollen allergen Lol p 2-A | 3.827971796 | 0.011998364 |
| Os04g0317800 | pollen allergen Phl p 2 | 3.499131819 | 0.014078969 |
| Os06g0556600 | pollen allergen Phl p 11 | 3.138908062 | 0.003136535 |
| Os12g0546800 | expansin-A26 | 3.138908062 | 0.008991554 |
| Os03g0106800 | expansin-B10 | 6.221623189 | 0.011550525 |
| Os03g0106900 | expansin-B1-like | 5.224194664 | 0.027081247 |
| Os08g0561900 | expansin-A32 | 4.95419631 | 0.03397062 |
| Os10g0548600 | expansin-B9 | 3.11849006 | 0.001361176 |
| Os03g0212800 | β-glucosidase 6 isoform X1 | 1.032586947 | 9.75E-16 |
| Os02g0131400 | β-glucosidase precursor | 1.436366009 | 0.004004821 |
| Os10g0323500 | β-glucosidase 34 | 1.815759811 | 0.0002305 |
| Os12g0420100 | β-glucosidase 38 | 2.154414422 | 0.0000689 |

**Supplementary Table S2.**
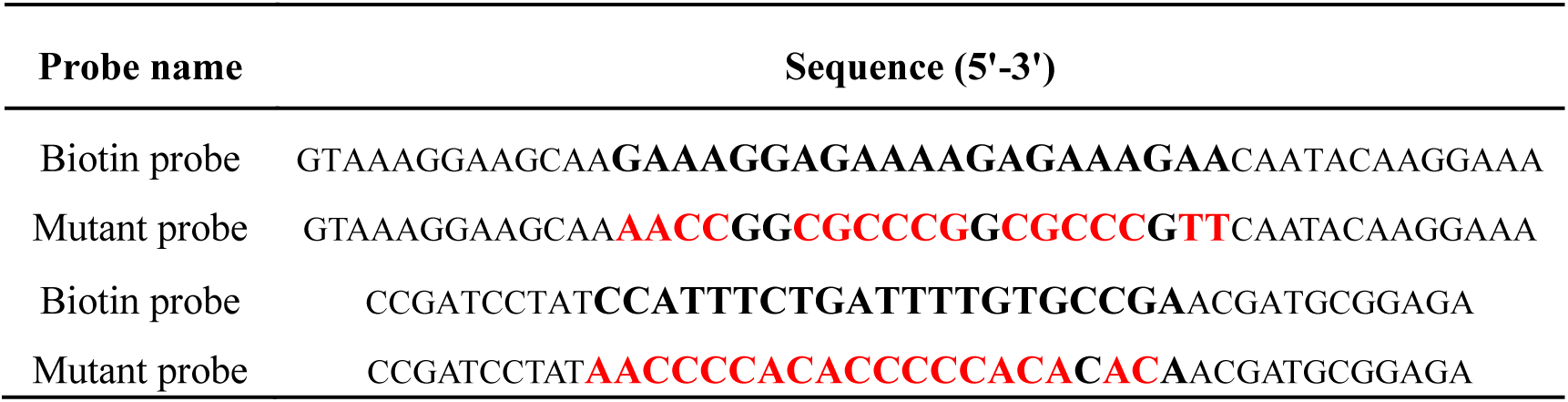
Probes used in EMSA.

**Supplementary Table S3.**
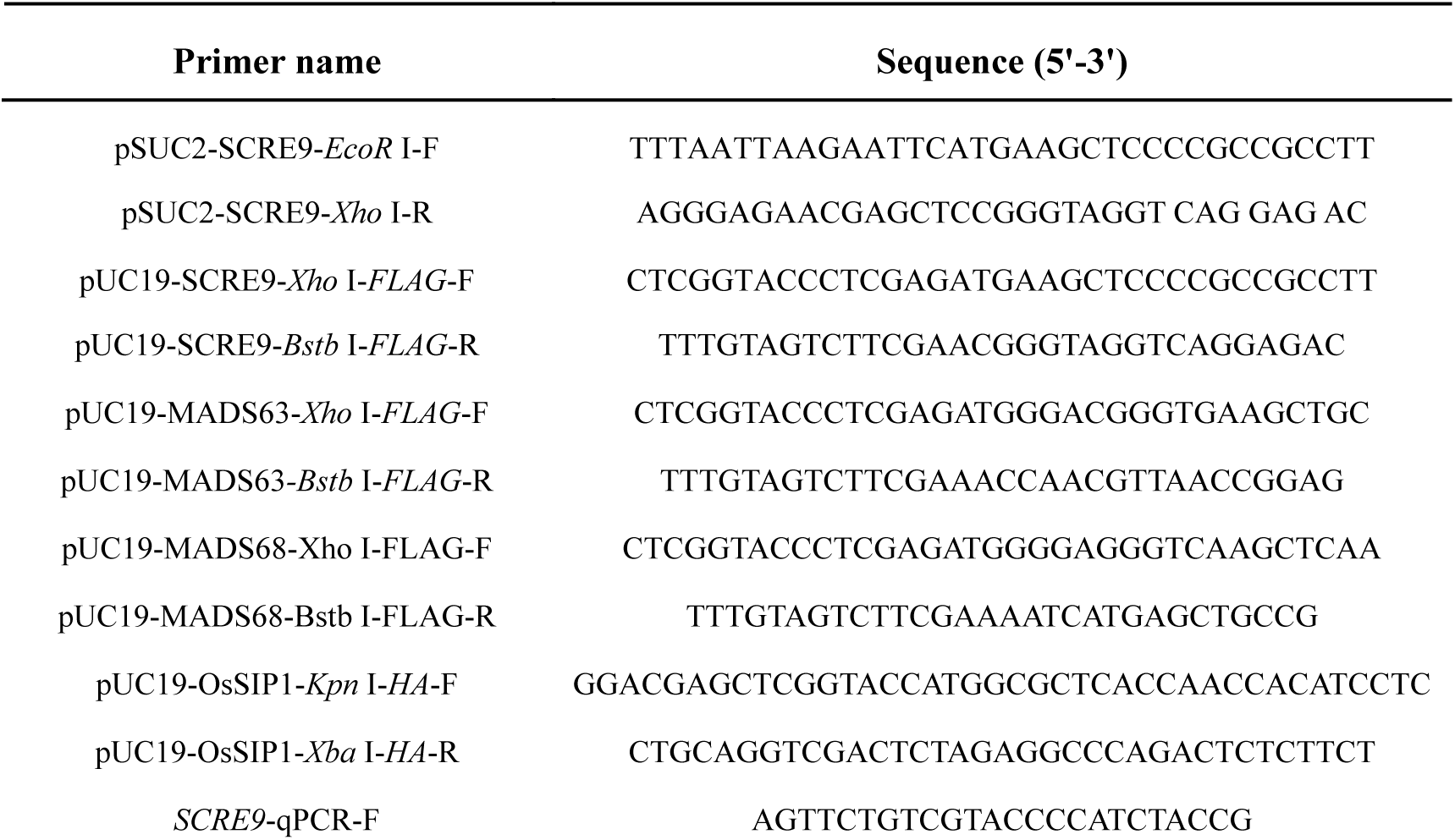
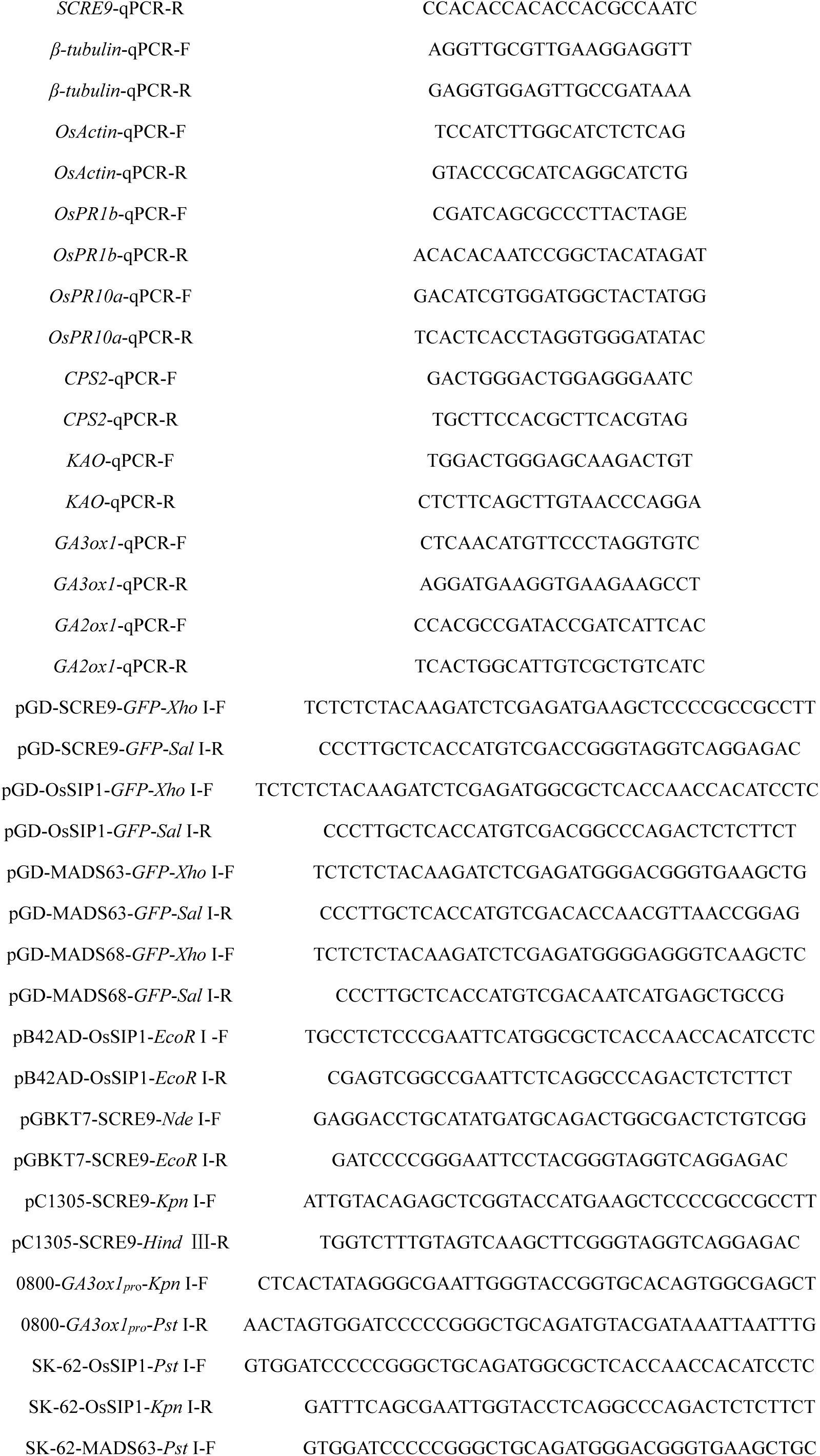
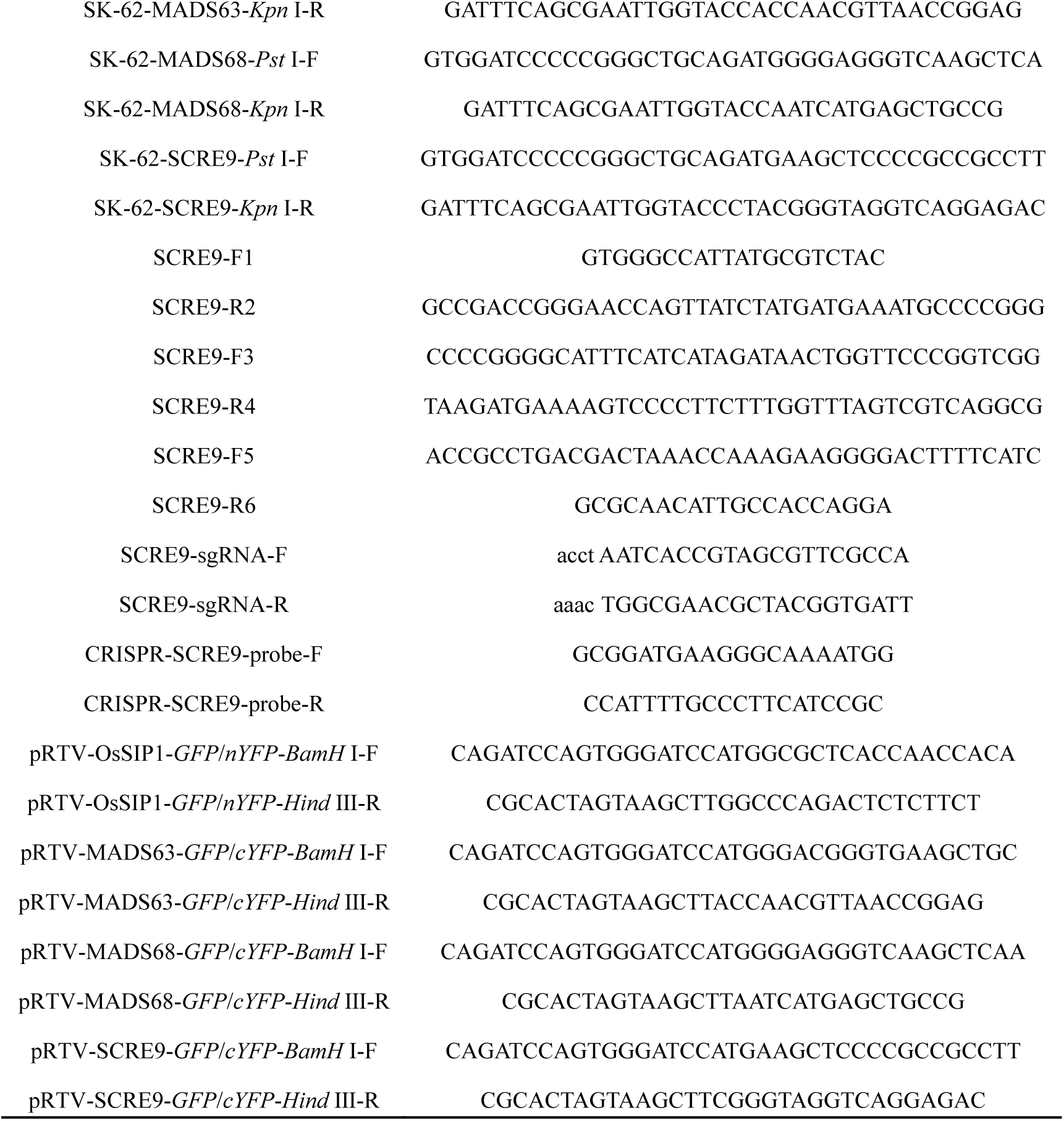
Primers used in this study.

